# The Low Complexity Domain of the FUS RNA Binding Protein Self-assembles via the Mutually Exclusive Use of Two Distinct Cross-β Cores

**DOI:** 10.1101/2021.08.05.455316

**Authors:** Masato Kato, Steven L. McKnight

**Affiliations:** Department of Biochemistry, UT Southwestern Medical Center, 5323 Harry Hines Boulevard Dallas, Texas 75390; Institute for Quantum Life Science, National Institutes for Quantum and Radiological Science and Technology, 4-9-1, Anagawa, Inage-ku, Chiba, Japan, 263-8555

## Abstract

The low complexity (LC) domain of the fused in sarcoma (FUS) RNA binding protein self-associates in a manner causing phase separation from an aqueous environment. Incubation of the FUS LC domain under physiologically normal conditions of salt and pH leads to rapid formation of liquid-like droplets that mature into a gel-like state. Both examples of phase separation have enabled reductionist biochemical assays allowing discovery of an N-terminal region of 56 residues that assembles into a labile, cross-β structure. Here we provide evidence of a non-overlapping, C-terminal region of the FUS LC domain that also forms specific cross-β interactions. We propose that biologic function of the FUS LC domain may operate via the mutually exclusive use of these N- and C-terminal cross-β cores. Neurodegenerative disease-causing mutations in the FUS LC domain are shown to imbalance the two cross-β cores, offering an unanticipated concept of LC domain function and dysfunction.

**Significance Statement:** Single amino acid changes causative of neurologic disease often map to the cross-β forming regions of low complexity (LC) domains. All such mutations studied to date lead to enhanced avidity of cross-β interactions. The LC domain of the fused in sarcoma (FUS) RNA binding protein contains three different regions that are capable of forming labile cross-β interactions. Here we describe the perplexing effect of amyotrophic lateral sclerosis (ALS)-causing mutations localized to the LC domain of FUS to substantially weaken its ability to form one of its three cross-β interactions. An understanding of how these mutations abet uncontrolled polymerization of the FUS LC domain may represent an important clue as to how LC domains achieve their proper biological function.

## Introduction

Upwards of 20% of the proteomes of eukaryotic cells consists of polypeptide sequences that are of low complexity (LC) (1, 2). These atypical polypeptide domains are composed of but a subset of the 20 amino acids normally required for proteins to assume their distinct, three dimensional shapes. As studied in the monomeric state in biochemical preparations outside of cells, LC sequences remain unfolded. As such, they have been described as intrinsically disorder regions (IDR’s) of the proteome.

Unbiased proteomic studies employing thermal perturbation as a probe for structural order within cells have shown that a surprisingly large fraction of protein domains thought to be intrinsically disordered may instead function in states of labile, structural order (3). Numerous reports published over the past several years have also given evidence that LC domains can undergo phase separation upon purification and incubation in test tube assays (4, 5). Purified LC domains can self-associate in a manner causing them to partition out of aqueous solution by forming liquid-like droplets. Following incubation, these droplets solidify into a gel-like state. If the interactions driving phase separation of LC domains properly reflect their biologic function, this line of research may help reveal how these unusual protein domains actually work in living cells.

Studies of the fused in sarcoma (FUS) RNA binding protein offered an early example of the unusual behavior of phase separation by an LC domain (6, 7). Incubation of the purified FUS LC domain initially triggers the formation of an opalescent suspension composed of liquid-like droplets. Sustained incubation of the same preparation leads to a more stable, gel-like state. Electron microscopy and X-ray diffraction analysis revealed FUS hydrogels to be composed of uniform, amyloid-like polymers (7). Unlike pathogenic, prion-like amyloids, FUS polymers are labile to disassembly upon dilution. A molecular structure of FUS polymers has been resolved by the use of solid state NMR spectroscopy (8). Among the 214 residues constituting the FUS LC domain, polymers were found to form via the organization of in-register, cross-β interactions localized between residues 39 and 95.

Structural studies of FUS polymers revealed two differences from pathogenic amyloids, such as α-synuclein or polymers formed from the Aβ fragment long understood to form hyper-stable aggregates in Alzheimer’s disease patients. First, FUS polymers reproducibly form the same, monomorphic structure via an N-terminally localized cross-β core. By contrast, pathogenic fibrils formed from α-synuclein or Aβ can adopt any of a number of different, inordinately stable structures (9, 10). Second, the subunit interface holding α-synuclein or Aβ polymers together are replete with hydrophobic amino acids believed to contribute to extreme polymer stability. The molecular structure of FUS polymers revealed but a single hydrophobic amino acid, proline residue 73, within the 56 residues constituting the subunit interface (8). The paucity of hydrophobic residues at the subunit interface of FUS polymers may, at least in part, explain polymer lability.

Studies of the LC domain of the hnRNPA2 protein have yielded similar findings. The LC domain of hnRNPA2 becomes phase separated into liquid-like droplets that, with time, also mature into a gel-like state (11, 12). Cross-β interactions formed by a region 40-50 residues in length also define hnRNPA2 LC domain polymers (11, 13, 14). Human genetic studies of patients suffering from various forms of neurological disease have identified mutations in the genes encoding hnRNPA1, hnRNPA2 and hnRNPDL (15, 16). These recurrent, disease-causing mutations commonly alter a conserved aspartic acid residue located within the cross-β core known to hold hnRNPA2 LC domain polymers together (11, 13, 17).

A simplistic interpretation of these observations offers that evolution has favored the presence of an aspartic acid residue within the cross-β cores formed from hnRNP LC domains as a means of tuning the balance of polymer stability/lability. Proximal disposition of this aspartic acid residue, as dictated by the in-register organization of the polymer interface, has been hypothesized to impart instability resulting from repulsive charge:charge interactions (13). Mutations removing these destabilizing interactions, especially when replacing the conserved aspartic acid residue with valine, may enhance polymer stability and lead to disease pathophysiology.

The relatively straightforward understanding of disease-causing mutations in the LC domains of three different hnRNP proteins does not translate to human genetic studies of FUS. Two of the most prominent, ALS-causing mutations within the FUS LC domain include a missense mutation changing glycine residue 156 to glutamic acid, and the deletion of glycine residues 174 and 175 (18–20). Neither of these mutations is anywhere close to the cross-β core of FUS that is located between residues 39 to 95.

The inability of our current understanding of the FUS cross-β core to explain substantive human genetic studies prompted re-examination of the pathway by which soluble monomers are captured by hydrogel samples composed of the intact LC domain of FUS. As will be described, these studies reveal surprising and unanticipated observations. The baroque pathway of subunit recruitment into existing FUS polymers offers an unconventional perspective as to how disease-causing mutations in the FUS LC domain manifest their pathophysiology at a molecular level. These studies further offer a means of understanding how the FUS LC domain is prevented from run-away polymerization via either of its cross-β cores.

## Results

### Identification of a region of the FUS LC domain critical for hydrogel binding

The sequence of the FUS LC domain is quasi-repetitive, containing 27 repeats of the tri-peptide sequence G/S-Y-G/S (Figure 1B). Early mutational studies gave evidence that the tyrosine residues of these repeats are correlatively important for the binding of FUS to hydrogel preparations formed from the LC domain of FUS itself, as well as its recruitment to RNA granules in living cells (7). In these experiments, hydrogels were prepared from a fusion protein linking mCherry to the native LC domain of FUS, and test proteins were GFP fusions to either the native LC domain of FUS, or mutated variants thereof. We have repeated these same experiments using 27 variants of the FUS LC domain bearing single tyrosine-to-serine mutations. Little or small effect on hydrogel binding was observed for 24 of the 27 variants. Three variants, wherein tyrosine residues 155, 161 or 177 were individually changed to serine, bound to mCherry:FUS hydrogels considerably less well than the native protein (Figures 1A and 1C). It is notable that these mutation-sensitive tyrosine residues localize far distal to the cross-β core housed between residues 39 and 95.

**Figure 1.**
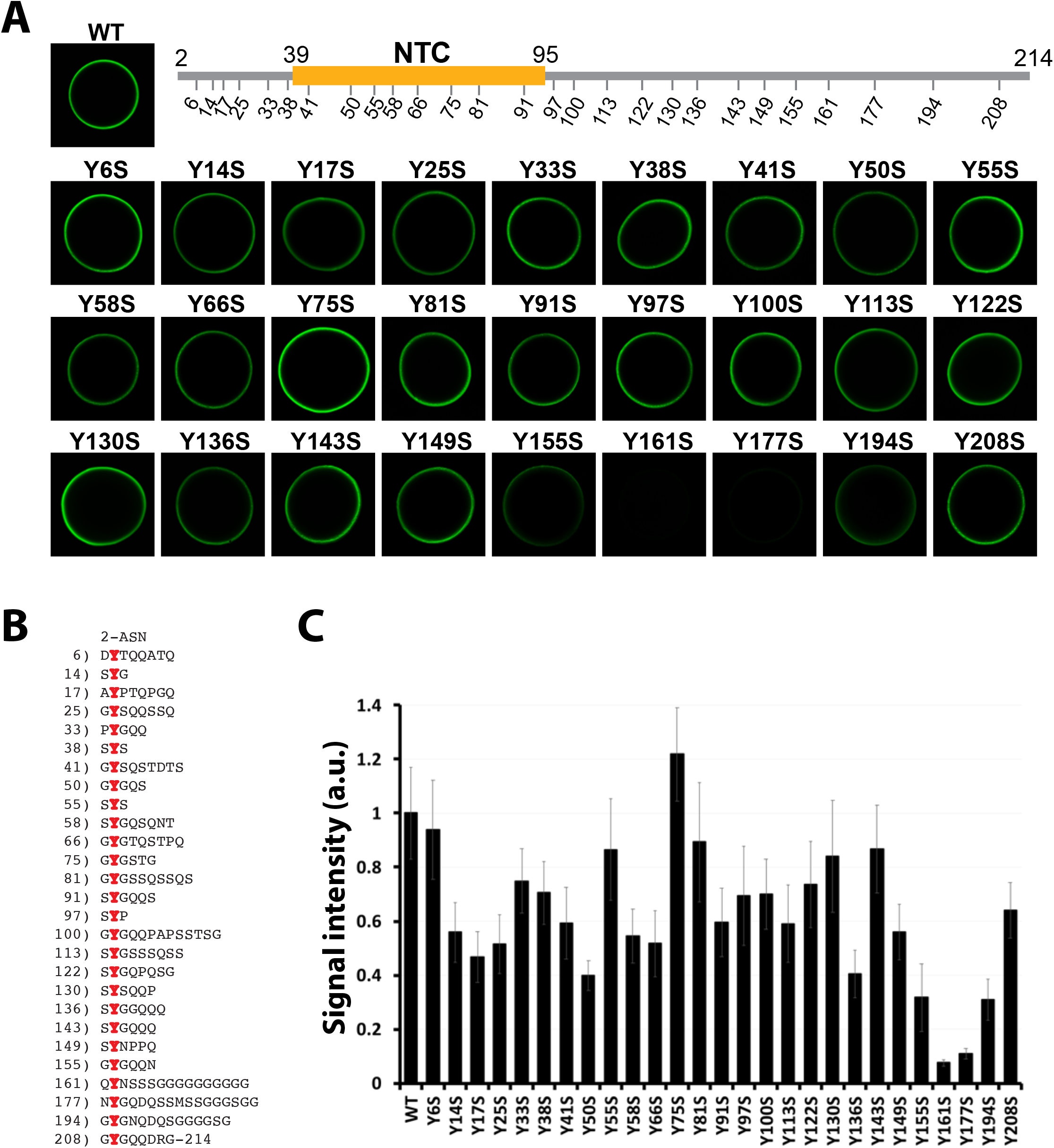
Identification of a C-terminal region of the FUS LC domain require for hydrogel binding. (A) Twenty seven individual tyrosine-to-serine mutations were made across the FUS low complexity domain. Yellow region of schematic diagram of the FUS LC domain corresponds to the location of the N-terminal cross-β structural core (NTC) determined by ssNMR (8). Native protein (WT) and individual point mutants were linked to GFP, expressed in bacteria, purified and tested in binding assays using hydrogel droplets formed from mCherry linked to the intact LC domain of FUS. Hydrogel binding activity was reduced most significantly by the Y161S and Y177S mutants. (B) Amino acid sequence of the FUS LC domain with tyrosine residues shown in red. Residue numbers for the Individual tyrosines are shown left. (C) Quantitation of hydrogel binding assays shown in panel A. Intensities are an average of three measurements.

Three sets of deletion mutants were prepared to further investigate regions of the FUS LC domain important for hydrogel binding. One set of deletions systematically truncated the LC domain from its C-terminus. As shown in Figure 2A, removal of 17 residues yielded a protein (ΔC1) that retained strong hydrogel binding activity. The next variant missing 34 residues (ΔC2) retained residual binding, yet all remaining deletion mutants were unable to bind hydrogels formed by the intact LC domain of FUS (Figure 2A). From these studies we define a boundary around residue 190 that marks the C-terminal location of a region required for monomeric GFP test proteins to bind mCherry hydrogels formed by the intact FUS LC domain

**Figure 2.**
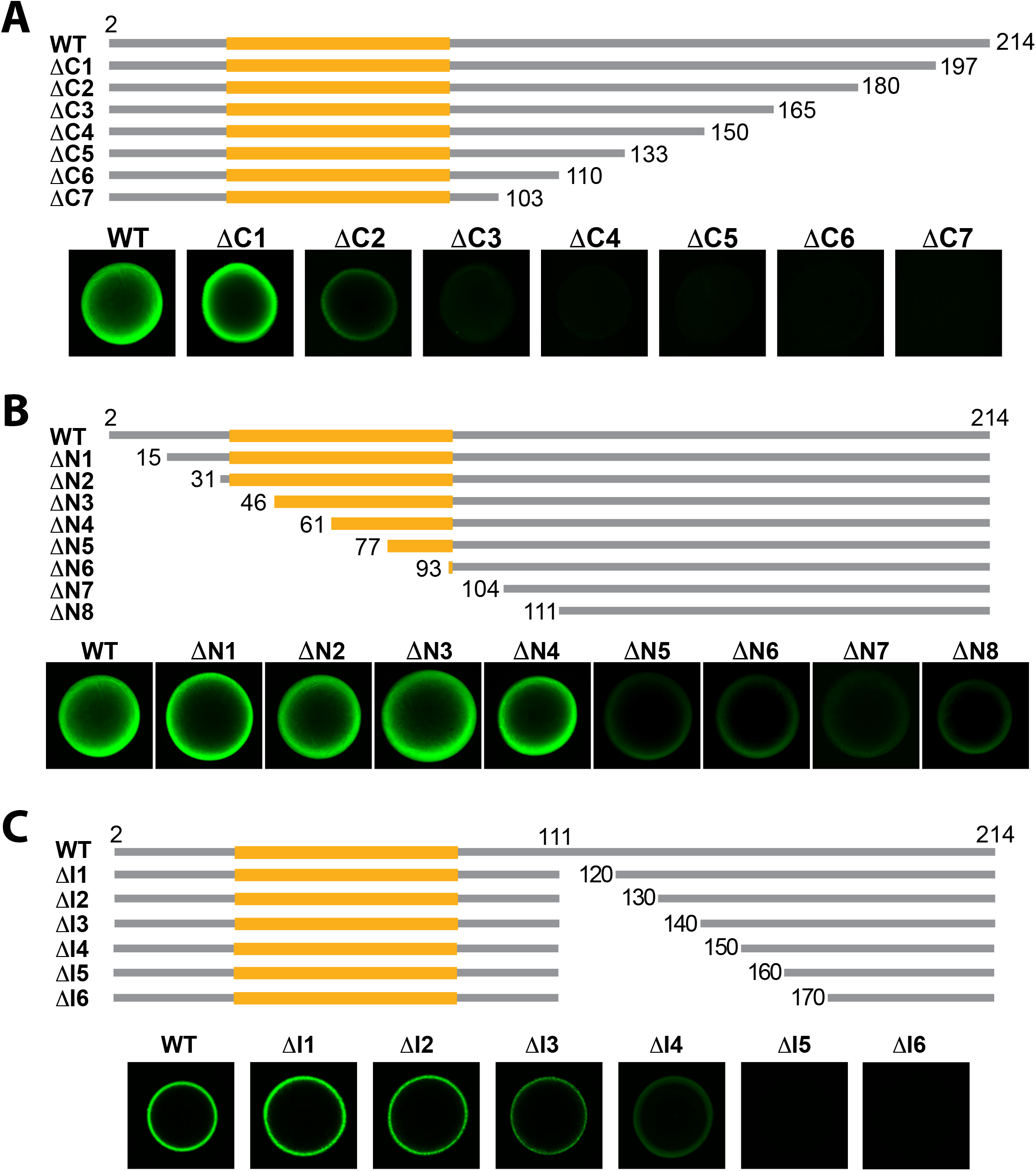
Hydrogel binding analysis of three sets of deletion variants of the FUS LC domain. (A) Seven deletion mutants progressing from the C-terminus of the FUS LC domain were linked to GFP, expressed in bacteria, purified and tested in hydrogel binding assays as described in Figure 1. One deletion mutant (ΔC1) displayed robust hydrogel binding activity, another (ΔC2) retained residual binding activity, and all other deletion mutants lost hydrogel binding activity. (B) Eight deletion mutants progressing from the N-terminus of the FUS LC domain were linked to GFP, expressed in bacteria, purified and assayed for hydrogel binding. Four deletion mutants (ΔN1-ΔN4) displayed robust hydrogel binding activity, and four deletion mutants (ΔN5-ΔN8) retained attenuated hydrogel binding activity. (C) Six internal deletion mutants progressing in ten amino acid increments from residue 111 were linked to GFP, expressed in bacteria, purified and assayed for hydrogel binding. Four internal deletion mutants (ΔI1-ΔI4) displayed hydrogel binding activity, and two did not (ΔI5, ΔI6). Yellow regions in schematic diagrams designate the location of the N-terminal cross-β core (NTC) of the FUS LC domain as defined by its atomic structure spanning residues 39-95 (8).

Analysis of N-terminal truncations of the FUS LC domain yielded the surprising observation that upwards of half of the region specifying the cross-β core, spanning residues 39-95, could be deleted without significantly affecting hydrogel binding (Figure 2B). The ΔN4 variant displayed strong hydrogel binding despite lacking 22 residues of the cross-β core. It was likewise surprising that four deletion mutants missing even greater amounts of the FUS LC domain, two of which eliminated the entire cross-β core (ΔN7 and ΔN8), retained attenuated but readily detectible hydrogel binding activity. In combination, these systematic mutagenesis experiments give evidence that a region of the FUS LC domain located far distal to the cross-β core is required for an unstructured test protein to bind hydrogel preparations of the FUS LC domain.

In a third set of deletion mutations, we initiated truncation downstream of the N-terminal cross-β core starting at residue 111. Deletions were extended from this point and enlarged at ten amino acid increments, internally removing as little as ten residues and extending up to as much as 60 residues. Each variant was expressed as a GFP fusion, purified and tested for binding to mCherry hydrogel samples formed from the intact FUS LC domain. The internal deletion missing 30 amino acids (ΔI3), and all variants missing fewer residues of the FUS LC domain, exhibited hydrogel binding. By contrast, the variant missing 40 internal residues exhibited reduced hydrogel binding, and those missing 50 or 60 residues revealed no binding (Figure 2C). In combination with studies of C-terminal truncations (Figure 2A), analysis of these internal deletion mutants define a region of roughly 40 amino acids, located between residues 150 and 190 of the FUS LC domain, required for monomeric test protein to bind mCherry:FUS hydrogels.

Interpretation of these experiments assumes that the mCherry:FUS hydrogel droplets are composed of polymers assembled via the same cross-β core whose structure was resolved at the atomic level by solid state NMR spectroscopy (8). Such studies were performed on the isolated LC domain of FUS not appended to mCherry or GFP. It was formally possible that polymers formed from the mCherry:FUS fusion protein used in this study might be different from polymers formed from the isolated FUS LC domain. To test this possibility we employed intein chemistry to ligate un-labeled GFP to a uniformly, ^13^C,^15^N-labeled form of the FUS LC domain. The latter protein was allowed to polymerize and evaluated by solid state NMR spectroscopy. As shown in Figure S1, polymers made from the segmentally labeled GFP:FUS chimeric protein yielded NMR spectra indistinguishable from polymers prepared from the isolated, uniformly labeled LC domain of FUS.

### Identification of a cross-β core within the C-terminal half of the FUS LC domain

The ΔN8 deletion variant contains residues 111-214 of the FUS LC domain. From here forward we designate this as the C-terminal half of the FUS LC domain. This protein fragment of 104 residues was expressed in bacterial cells, purified, and incubated under conditions receptive to phase transition (Materials and Methods). Upon incubation at neutral pH and physiological concentration of monovalent salt (Materials and Methods), this C-terminal half of the FUS LC domain became phase separated into a gel-like state. When observed by transmission electron microscopy, the hydrogel was found to be composed of uniform, unbranched polymers (Figure S2A). X-ray diffraction analysis of hydrogels formed from the C-terminal half of the FUS LC domain revealed diffractions rings at 4.7 and 10 Å (Figure S2B). Finally, when analyzed by semi-denaturing agarose gel electrophoresis (SDD-AGE), the observed polymers were labile to disassembly (Figure S2C).

Truncations of the C-terminal half of the FUS LC domain were prepared having N-termini at residues 141, 145, 150, 155 and 160, together with a common C-terminus at residue 214 (Figure S3A). Following incubation under conditions receptive to phase transition, each variant was evaluated for its capacity to polymerize both by time-dependent acquisition of thioflavin-T fluorescence and transmission electron microscopy (Figure S3B). All variants were observed to form homogenous polymers save for the most truncated fragment bearing an N-terminus at residue 160. We thus conclude that the region of the FUS LC domain located between residues 155 and 214 specifies a secondary cross-β core distinct from that of the N-terminal half of the FUS LC domain characterized extensively in previous studies (8).

The laboratory of Robert Tycko has recently described the structure of cross-β polymers formed from the C-terminal half of the FUS LC domain (21). The structural core of the Tycko polymers, whose atomic fold was resolved by cryo-electron microscopy, is specified by residues 112-150 of the FUS LC domain. This region of 39 amino acids is not required for soluble test protein to bind to mCherry hydrogels formed from the intact LC domain of FUS (Figure 2C). It is likewise clear that the cross-β core described by Tycko and colleagues is entirely distinct from the C-terminal cross-β core described herein. Truncated variants of the C-terminal half of the FUS LC domain completely lacking residues 111-150 are fully capable of polymerization (Figure S3B).

To compare cross-β polymers corresponding to those described by the Tycko lab with those described in this study, fragments of the FUS LC domain spanning either residues 111-160 or 141-214 were purified and allowed to polymerize under conditions receptive to phase separation. Both fragments readily formed cross-β polymers that were evaluated by electron microscopy, X-ray diffraction, semi-denaturing agarose gel electrophoresis and solid state NMR spectroscopy. Both samples yielded homogeneous, unbranched polymers as viewed by electron microscopy (Figure S4A). Both samples revealed X-ray diffraction images consistent with cross-β amyloid-like polymers (Figure S4B). Both samples of polymers were labile to disassembly when subjected to SDD-AGE (Figure S4C). Finally, the two samples revealed different spectra as evaluated by solid state NMR spectroscopy (Figure S4D). For reasons we do not yet understand, the region specifying the cross-β core characterized by Tycko and colleagues (residue 112-150) is not required for hydrogel binding by the intact FUS LC domain (Figures 1 and 2). By contrast, the region of the FUS LC domain specifying the C-terminal cross-β core described in this report (155-190) is vitally required for hydrogel binding.

From here forward we will refer to the region spanning residues 39-95 of the FUS LC domain as specifying the boundaries of the predominating, N-terminal cross-β core (NTC) of the FUS LC domain (8). We will further term the region spanning residues 155-190 as specifying the boundaries of the secondary, C-terminal cross-β core (CTC) of the FUS LC domain that has been characterized in the present study.

In order to investigate the possible relationship between this secondary, C-terminal cross-β core and the capacity of soluble FUS LC domain monomers to bind mCherry hydrogels formed from the intact LC domain of FUS (Figures 1 and 2), we investigated the effects of single tyrosine-to-serine (S-to-Y) mutations upon polymerization of the fragment of the FUS LC domain spanning residues 141 to 214. As shown in Figure 3, four of the single Y-to-S mutants (Y143S, Y149S, Y194S and Y208S) polymerized in a manner indistinguishable from the wild-type fragment. By contrast, three of the Y-to-S mutants (Y155S, Y161S and Y177S) were severely compromised in polymerization capacity (Figure 3). This pattern of effects on polymerization closely mimics the pattern of effects of the same seven Y-to-S mutants on hydrogel binding by the otherwise intact LC domain of FUS (Figure 1).

**Figure 3.**
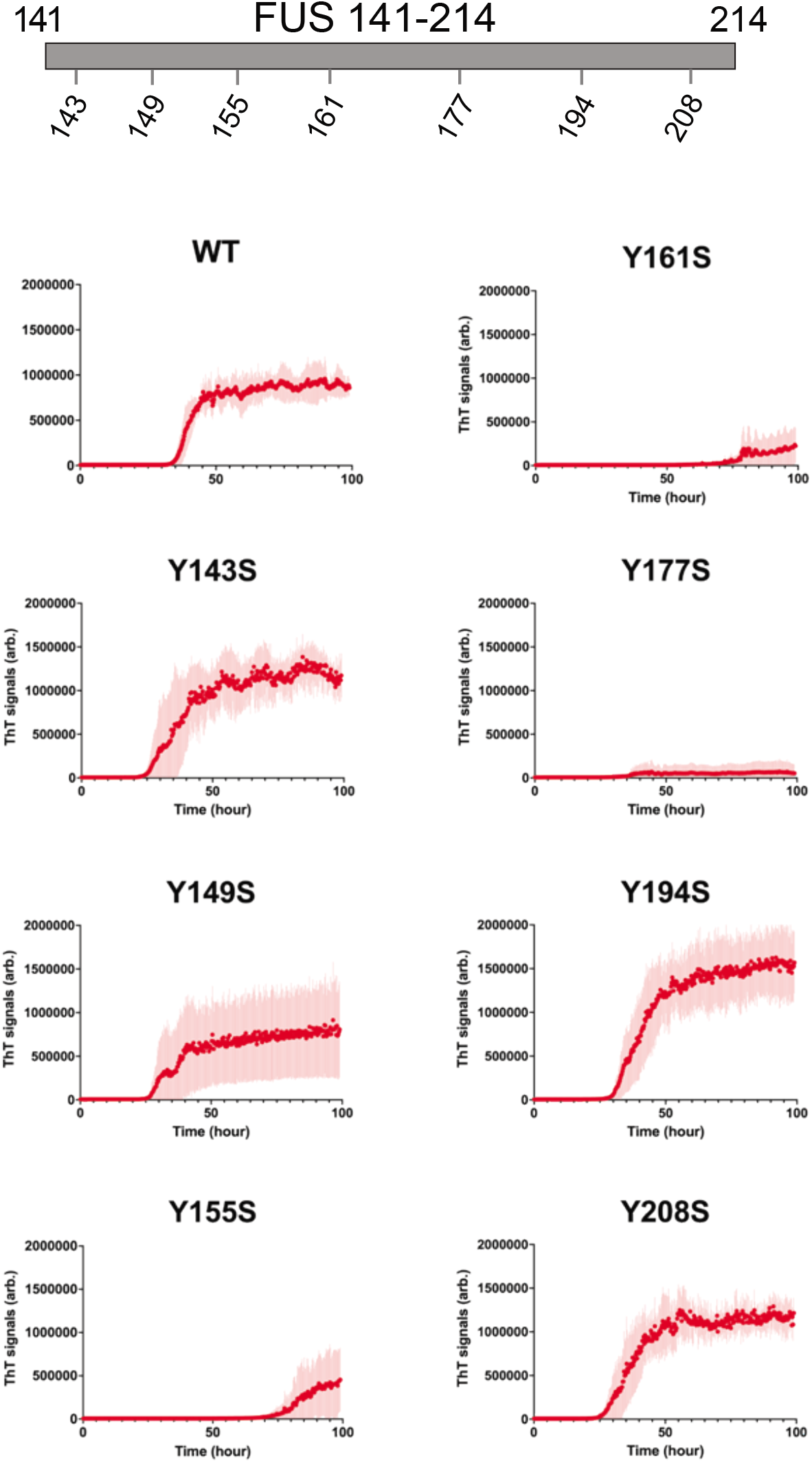
Polymerization capacity of a fragment of the FUS LC domain spanning residues 141-214. Acquisition of thioflavin-T fluorescence as a function of time was compared for fragments of the FUS LC domain spanning residues 141-214 bearing either the native sequence or that of variants carrying a single tyrosine-to-serine mutation. Each graph displays fluorescence increase (X-axis) relative to time of incubation (Y axis). Four mutants, including Y143S, Y149S, Y194S and Y208S, revealed evidence of polymerization similar to the wild type protein. Three mutants, including Y155S, Y161S and Y177S, revealed substantially impeded capacity for polymerization.

### Evidence of molecular specificity in elongation of both NTC and CTC cross-β polymers

In efforts to complete a more thorough analysis of the two regions allowing for self-association of the FUS LC domain, we prepared hydrogels composed of either the NTC (residues 2-110) or the CTC (residues 111-214) and tested their respective abilities to bind two GFP-tagged test proteins. Both hydrogels were made from mCherry fusion proteins. The two GFP test proteins used to interrogate the two hydrogel samples included one composed of the N-terminal half of the LC domain (residues 2-110), or a second composed of the C-terminal half of the LC domain (residues 141-214). The latter protein was purposely truncated at its N-terminus to remove the region shown recently by Tycko and colleagues to be capable of forming cross-β polymers (21).

As shown in Figure 4, the GFP fusion protein linked to the N-terminal half of the LC domain (residues 2-110) bound only the hydrogel made from the NTC itself. The GFP fusion protein linked to the CTC (residues 141-214) bound only the hydrogel made from the C-terminal half of the LC domain. Having observed that the GFP fusion protein containing the NTC does not bind to mCherry:CTC hydrogels, and that the GFP fusion protein containing the CTC does not bind to mCherry:NTC hydrogels, we conclude that these binding reactions offer evidence of molecular specificity.

**Figure 4.**
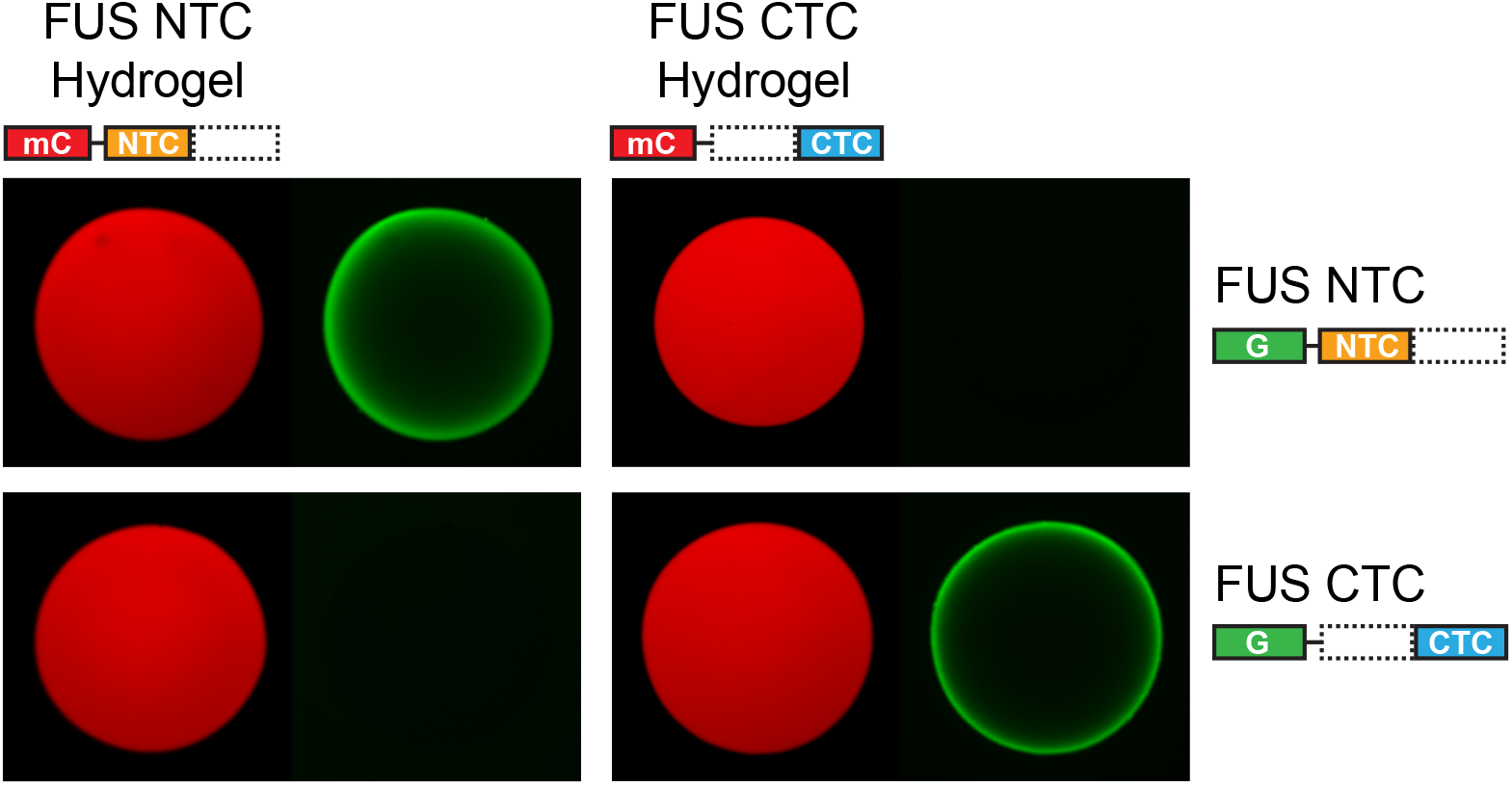
Specificity of binding of N- and C-terminal halves of the FUS low complexity domain to hydrogel polymers formed from the same regions. mCherry fusion proteins linked to either the N-terminal half (residues 2-110) or C-terminal half (residues 111-214) of the FUS LC domain were challenged with soluble GFP fusion proteins corresponding to the same two halves of the protein. mCherry hydrogels composed of the N-terminal half of the FUS LC domain bound the GFP fusion protein linked to the same, N-terminal half of the protein, but not the C-terminal half (left panels). mCherry hydrogels composed of the C-terminal half of the FUS LC domain bound the GFP fusion protein linked to the same, C-terminal half of the protein, but not the N-terminal half (right panels).

### Kinetic formation rates and stabilities of the N- and C-terminal cross-β cores of the FUS LC domain

Purified, tag-free fragments corresponding to the N- and C-terminal halves of the FUS LC domain (Figure 5A) were diluted out of denaturant and monitored for polymerization by acquisition of thioflavin-T fluorescence. As shown in Figure 5B, the fragment corresponding to the C-terminal half of the FUS LC domain polymerized more rapidly than the fragment corresponding to the N-terminal half. In order to measure the stabilities of the NTC and CTC polymers, assembled polymers were exposed to graded increases in temperature. Release of soluble monomers was monitored by reverse-phase column chromatography (Figure 5C). Roughly, half of the CTC polymers were soluble at 50°C, and all of the sample was converted to monomers at 60°C. By contrast, the NTC polymers required 10°C higher temperature to achieve, respectively, half-maximal or full disassembly.

**Figure 5.**
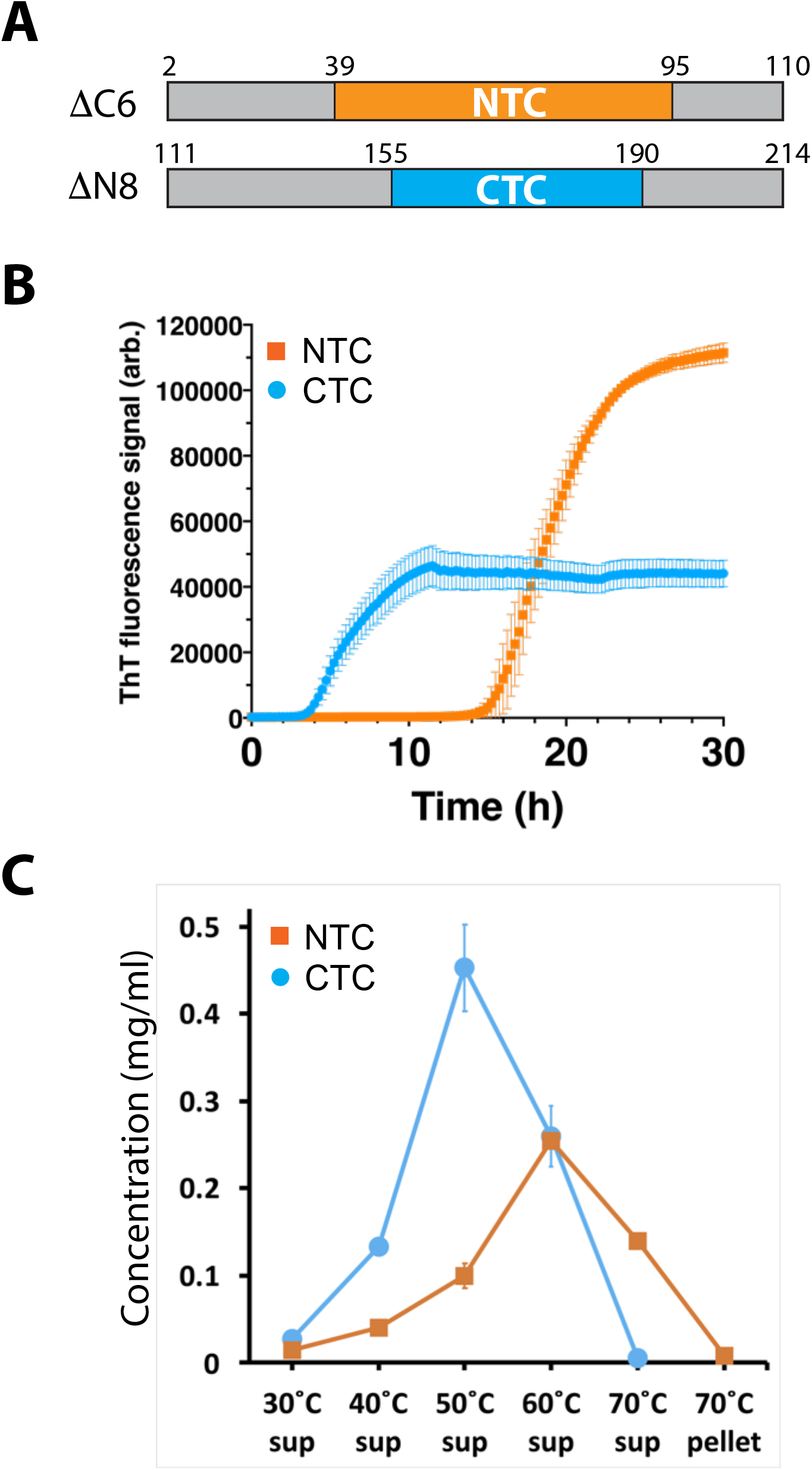
Measurements of polymerization rates and stabilities of polymers formed from the N- and C-terminal halves of the FUS low complexity domain. (A) Schematic diagram of protein fragments corresponding to N- and C-terminal halves of the FUS low complexity domain. Boxed region shown in yellow corresponds to structural boundaries of the N-terminal cross-β core (NTC) of the FUS LC domain (residues 39-95). Boxed region shown in cyan corresponds to functional boundaries of the C-terminal cross-β core (CTC) forming region of the FUS LC domain (residues 155-180). (B) Proteins fragments shown in panel A were expressed in bacteria, purified and incubated under conditions of neutral pH and physiological monovalent salt for 30 hrs. Thioflavin-T fluorescence (Y axis) was measured as a function of time (X axis). (C) Polymerized samples of NTC and CTC polymers were incubated at varying temperatures (X axis) and monitored for the release of soluble monomers by reverse-phase column chromatography (Y axis).

### ALS-causing variants destabilize the C-terminal cross-β core of the FUS low complexity domain

Variants of the FUS LC domain reported to pre-dispose patients to amyotrophic lateral sclerosis (ALS) include a glycine-to-glutamic acid missense mutation of residue 156 (G156E) (18) and the deletion of glycine residues 174 and 175 (ΔG174/G175) (20, 22, 23). Recognizing that these variants map within the C-terminal cross-β core of the FUS LC domain, we initially expressed and purified both variants as GFP fusion proteins in the context of the isolated C-terminal core. Upon incubation under conditions normally leading to phase transition, we were surprised to observe that neither variant was able to form cross-β polymers within a time frame in which the native protein readily polymerized.

In order to investigate these observations more carefully, purified, tag-free monomeric protein was tested for polymerization via assays of thioflavin-T fluorescence. Figure 6B shows polymerization assays for the truncated C-terminal core (FUS 141-214) in its native sequence configuration as compared with variants carrying either the G156E missense mutation or G174/G175 deletion (ΔG174/G175) (Figure 6A). Although anticipating that the ALS-disposing variants might prompt the formation of aberrantly stable or more rapidly forming cross-β polymers, we observed no detectable polymerization for the ΔG174/G175 variant and significantly delayed polymerization for the G156E variant (Figure 6B). When tested in binding assays to hydrogels formed from the C-terminal cross-β core (CTC) alone, weak binding (G156E) or no binding (ΔG174/G175) was observed for these ALS-disposing mutants (Figure 6C).

**Figure 6.**
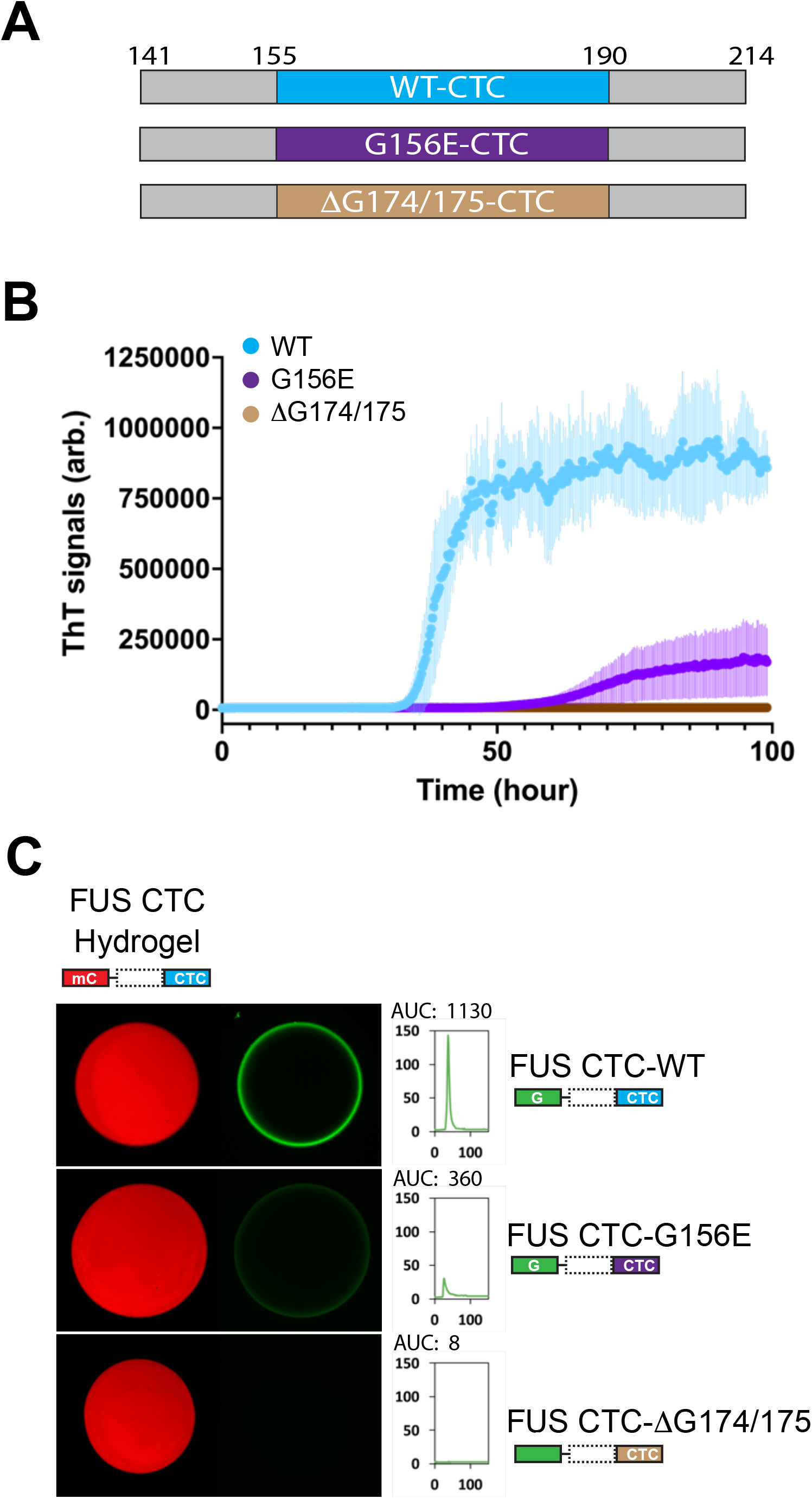
Measurements of polymerization and hydrogel binding of ALS-causing mutations as assayed within the isolated C-terminal cross-β core of the FUS low complexity domain. (A) Schematic diagram of protein fragments used to express native FUS LC domain (cyan) and ALS-causing variants (G156E = purple; ΔG174/175 = tan). (B) The region spanning residues 141-214 of the FUS LC domain was expressed in its native form as well as when carrying the G156E or ΔG174/G175 ALS-disposing variants. Purified protein was incubated under conditions of neutral pH and physiological monovalent salt for 100 hours. Thioflavin-T fluorescence (Y axis) was measured as a function of time (X-axis), giving evidence of polymerization by the native CTC fragment (blue line), delayed polymerization for the G156E variant (purple line), and no polymerization for the ΔG174/G175 variant (tan line). (C) The region spanning residues 111-214 of the FUS LC domain containing the C-terminal cross-β core was expressed as a GFP fusion in its native form as well as when carrying the G156E or ΔG174/175 ALS-disposing variants. Purified GFP-tagged proteins were incubated with hydrogels formed from mCherry linked to the C-terminal half of the FUS LC domain. Hydrogel binding activity evident for the GFP fusion linked to the native C-terminal half of the FUS LC domain was attenuated for the G156E variant, and absent for the ΔG174/G175 variant.

The G156E or ΔG174/G175 ALS-disposing variants were further analyzed in the context of the full-length LC domain of FUS by two assays: (i) acquisition of thioflavin-T fluorescence; and (ii) hydrogel binding. For the former assay, tag-free proteins corresponding to the native LC domain of FUS, and both ALS-disposing variants, were incubated under conditions of neutral pH and physiologic monovalent salt in the presence of thioflavin T. As shown in Figure 7A, both ALS-disposing variants acquired thioflavin-T fluorescence considerably more rapidly than the native FUS LC domain. This result, which is consistent with published studies of the G156E ALS-disposing variant of the FUS LC domain (24, 25), was notable in revealing the opposite pattern of polymerization from that observed when the three proteins were studied in the context of the isolated, C-terminal half of the FUS LC domain (Figure 6B).

**Figure 7.**
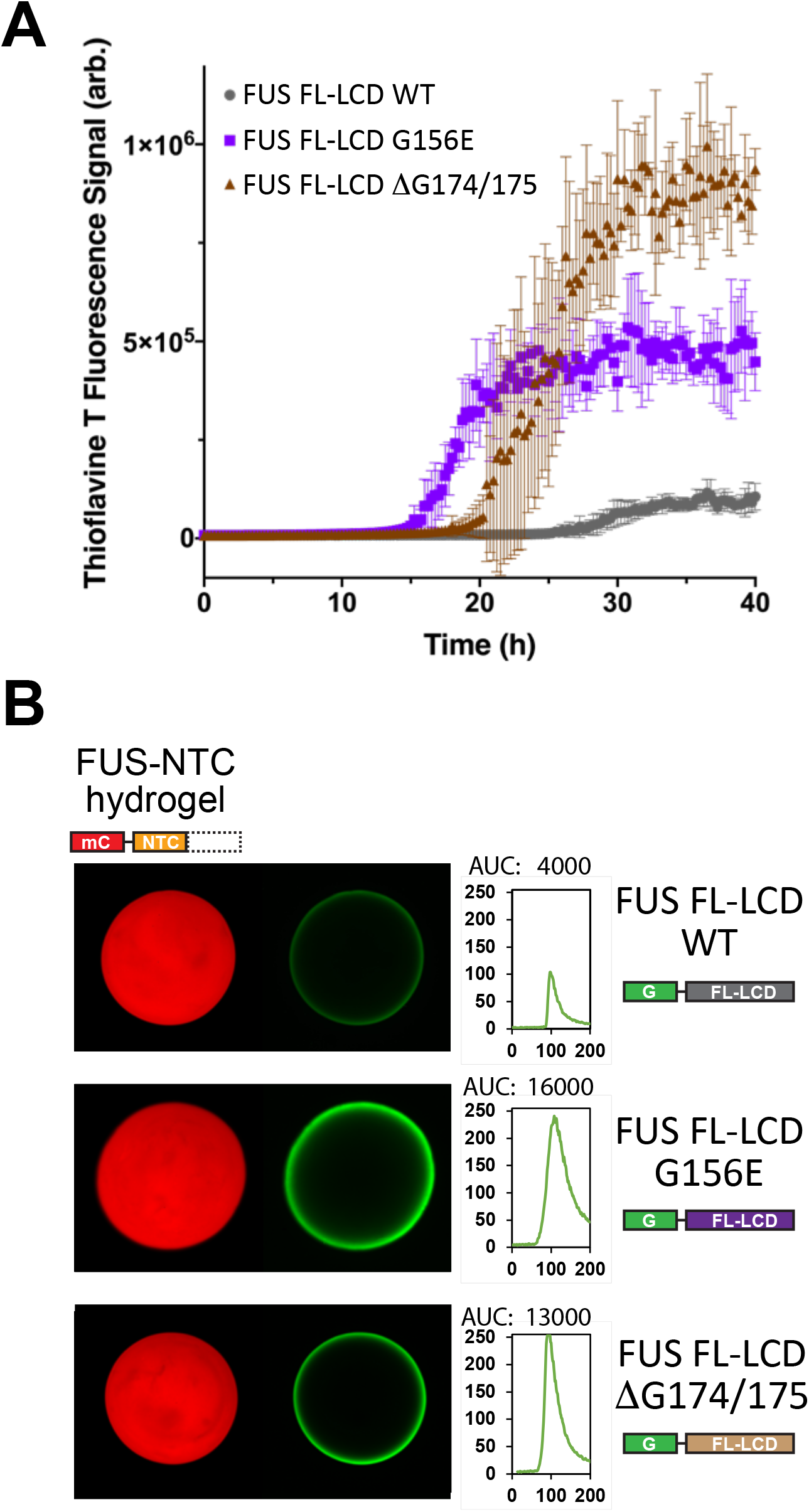
Measurements of polymerization and hydrogel binding of ALS-causing mutations as assayed within the intact low complexity domain of FUS. (A) Full length derivatives of the FUS low complexity domain bearing the native amino acid sequence (WT), or that carrying either of two ALS-disposing variants, were expressed as 6His-tagged proteins in bacteria, purified and incubated under conditions of neutral pH and physiological monovalent salt in the presence of thioflavin-T. Y axis presents thioflavin-T fluorescence, X-axis presents time of incubation. (B) The same three segments of the FUS low complexity domain were expressed as GFP-tagged proteins in bacteria, purified, and incubated under conditions of neutral pH and physiological monovalent salt with hydrogels formed from mCherry linked to the isolated N-terminal cross-β core of the FUS low complexity domain. Scans depicted to right of hydrogel images present quantitation of GFP signal intensity as measured at hydrogel perimeters. The G156E and ΔG174/G175 variants of the FUS low complexity domain (bottom two rows) displayed between 3- and 4-fold greater binding intensities than that observed for the native FUS protein (top row).

For hydrogel binding assays, each of the three proteins was linked to GFP, expressed in bacteria, purified and incubated with mCherry hydrogels formed from the N-terminal half of the FUS LC domain. To our surprise, both ALS-disposing variants bound more prominently to mCherry hydrogels formed from the N-terminal cross-β core (NTC) alone (Figure 7B). Why do variants carrying mutations that inactivate the C-terminal cross-β core bind NTC hydrogels more strongly than the intact LC domain of FUS? This observation, we propose, may offer an unanticipated clue as to how low complexity domains function in living cells.

## Discussion

In initiating the experiments described herein, we anticipated a simple pathway for the binding of soluble FUS monomers to pre-cast hydrogels. Given the presence of the structured N-terminal core (NTC) as the defining feature of mCherry hydrogels formed by the intact LC domain of FUS, our expectation was that the soluble, unstructured test protein would be capable of sampling the contours of the folded NTC located at polymer termini and simply slot itself into the existing protein fold. If so, the amino acid region specifying the NTC would have been the most important region of the FUS LC domain required for a soluble test protein to bind to hydrogel droplets. This expectation was not met. We instead observed that a distinct, C-terminal region of the FUS LC domain was considerably more important for hydrogel binding. These unexpected observations led to characterization of a different and non-overlapping cross-β forming region that we now designate as the C-terminal core (CTC) of the FUS LC domain.

What are we to make of these unexpected observations? What follows are hypothetical thoughts supported only in part by experimental observations. We offer these thoughts as correlates that may be helpful in considering how the FUS LC domain might achieve its biological function in a manner avoiding run-away polymerization from either the NTC or CTC.

### Correlate 1 – cross-β formation is achieved more readily by two unstructured regions than if one region already exists in a cross-β conformation

The sole way in which the FUS LC domain can exist in the unique cross-β structure that has been resolved by solid state NMR spectroscopy is if at least two molecules have coalesced. In other words, the cross-β conformation cannot be assumed by a monomer on its own. Once two molecules have assembled into the cross-β state, conformational freedom of the amino acids localized within the structured region is restricted. If one copy of the FUS LC domain exists in the structurally ordered state as the terminal cap of an existing cross-β polymer, and another is free and unstructured, we hypothesize the latter to have difficulty in finding the former.

It has been extensively postulated that π:π stacking of aromatic amino acid side chains may be important for self-associative interactions between unstructured LC domains (26). The FUS LC domain contains 27 tyrosine residues that are important for self-association of the protein (Figure 1) (7). Once assembled into the cross-β structural state, a subset of these tyrosine residues become conformationally restricted (8). We hypothesize that conformational restriction may impede the ability of an unstructured monomer to participate in tyrosine:tyrosine-mediated π:π stacking interactions within the region imposed to exist in the structurally order state.

If correct, this correlate may explain why the C-terminal region of FUS monomers is more important than the N-terminal region for a soluble test protein to bind hydrogels composed of NTC polymers. The LC domain of the soluble, GFP-tagged monomers whose binding is being assayed exists in an unstructured state. Likewise, the C-terminal region of FUS subunits already existing in hydrogel polymers is also unstructured so long as the polymers are assembled by the NTC (8). Since the C-terminal regions of both the hydrogel recipient of binding and test monomers are unstructured, we predict that the probability of forming a CTC cross-β interaction should exceed that of an NTC interaction.

### Correlate 2 – formation of both NTC and CTC cross-β structures within a single polypeptide is sufficiently unfavorable to impart mutual exclusion

Hydrogels formed by the intact LC domain of FUS contain uniform polymers thousands of subunits in length. These FUS polymers are held together by cross-β interactions specified by the NTC (8). The unstructured C-terminal region of the FUS LC domains extends laterally from polymers in bottle brush fashion, such that each polymer displays thousands of free and unstructured C-terminal regions. Here we show that when challenged with a soluble, unstructured test monomer, binding by the test protein is considerably more dependent upon the sequences of its C-terminal region than its N-terminal region.

Each polymer within a hydrogel has but two termini exposing the molded conformation of the structured, NTC cross-β assembly. By contrast, the same polymers display thousands of laterally extending C-terminal regions that are unstructured. Given this fact, coupled with correlate 1 as described above, it comes as no surprise that binding of the unstructured test protein is inordinately reliant upon the integrity of its C-terminal domain (Figures 1 and 2).

These findings are at odds, however, with simple microscopic observations of the process of polymer growth (7). Following co-incubation of mCherry:FUS polymers with soluble GFP:FUS test monomers, we used TIRF microscopy to image polymer growth as a function of time. As shown in Figure 8A, we observed no evidence of GFP binding to the lateral sides of pre-assembled mCherry:FUS polymers. Instead, time-dependent polymer growth revealed GFP-labeled termini. From such studies we offer that C-terminal interactions between the soluble test protein and the lateral sides of existing polymers must be unstable. It is possible that transient π:π stacking interactions via tyrosine side chains may enable these unstable, CTC:CTC interactions.

**Figure 8.**
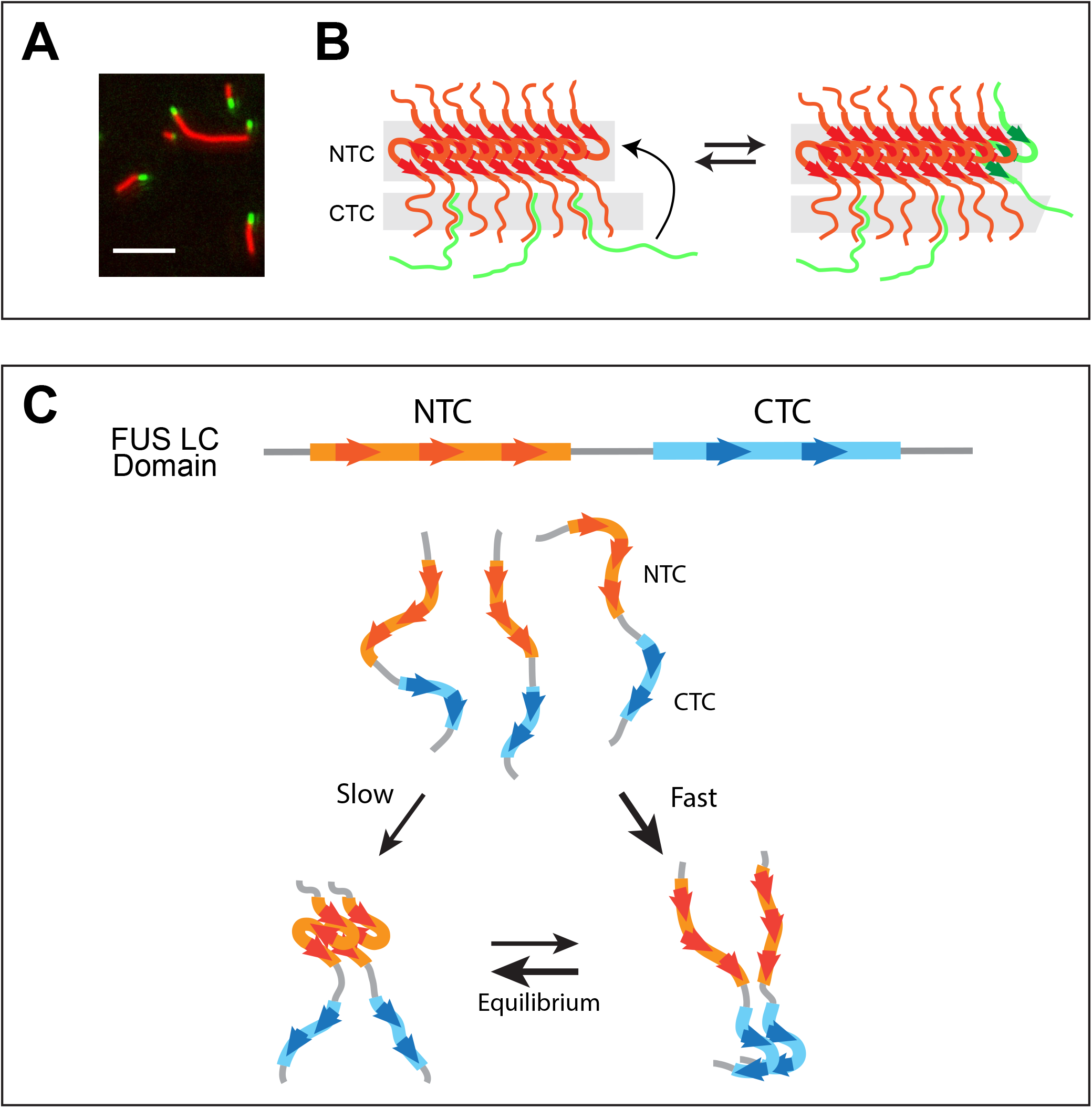
Hypothetical pathway of FUS low complexity domain binding to hydrogel polymers and conceptual pathway for initial self-association of soluble monomers. (A) TIRF microscopic imaging of mCherry:FUS polymers incubated with GFP:FUS monomers. Copolymerization of GFP test protein into existing mCherry:FUS polymers can be seen as green tips extending from red fibrils. (B) Schematic conceptualization of process of co-polymerization. Existing FUS polymer (red) is interpreted to initially bind soluble FUS monomers (green) via interactions wherein the CTC of test proteins attempts to form cross-β interaction with lateral surfaces of existing polymers (left). Initial, unstable interaction is interpreted to be in equilibrium with co-polymerization onto the N-terminal cross-β core (right). (C) Soluble, unstructured FUS low complexity domain monomers (top) are interpreted to preferentially self-associate via C-terminal cross-β core interactions (right). Slower yet more stable N-terminal cross-β self-association (left) is proposed to isomerize from dimers held together by less stable, C-terminal cross-β core. Simultaneous formation of N- and C-terminal cross-β interactions within a single polypeptide is interpreted to be impermissible.

We speculatively attribute these discordant observations to the fact that, owing to the correlate of mutual exclusion, a stable CTC cross-β structure cannot assemble on subunits of a polymer formed by the NTC cross-β structure. We instead imagine – as schematized in Figure 8B - that transient attempts towards CTC formation weakly adhere the soluble GFP-labeled test protein to mCherry-labeled hydrogel polymers. Stable hydrogel binding is limited to the event wherein the test protein co-polymerizes onto termini of existing polymers. In other words, C-terminal interactions between test protein and hydrogel may weakly retain the test protein in a position sufficiently proximal to polymer termini to enable execution of an unfavorable molecular event. Even though, according to correlate 1, binding of an unstructured N-terminal region to the cross-β structure at polymer termini is disfavored, it eventually transpires.

A more obvious indication of mutual exclusivity derives from structural studies of hydrogel polymers formed from the intact LC domain of FUS. Solid state NMR studies of these polymers have resolved the monomorphic structure of the N-terminal cross-β core (8). The in-register conformation of protomers held together by the NTC cause the unstructured, C-terminal domain of FUS to protrude laterally from the polymer core in a specified geometry. These extending, C-terminal regions of the polymer are flexibly positioned 4.7 Å apart. That the aligned, C-terminal regions do not adopt the cross-β conformation readily observed when the isolated CTC is incubated at high concentration (Figures 3, 4 and S2) bolsters the concept that individual molecules of the FUS LC domain cannot simultaneously form both N- and C-terminal cross-β cores.

Finally, the concept of mutual exclusion may explain the perplexing data shown in Figure 7B. mCherry hydrogel droplets composed solely of the NTC of FUS were challenged with GFP-tagged protein corresponding to the full-length LC domain bearing its native sequence, or that of either ALS variant. We offer that the enhanced binding of the latter proteins reflects the fact that, unlike the native LC domain, they cannot form CTC cross-β structures. According to correlate 2, mutual exclusion predicts an impediment to co-polymerization with NTC polymers of the hydrogel if the test protein can itself form CTC cross-β interactions. Since the native FUS LC domain can form CTC interactions, but the ALS variants cannot, the latter test proteins are able to bind NTC-only hydrogels more readily than the former.

### Implications of correlates 1 and 2

Correlates 1 and 2 represent simplified interpretations of the data included in this and earlier studies of the FUS LC domain. Despite their hypothetical nature, these correlates may be useful in thinking about questions of both narrow focus and potentially broad significance.

On the more narrow side, these correlates allow us to consider how certain ALS-causing mutations within the FUS LC domain might lead to aberrant protein aggregation. We have found that the G156E and ΔG174/G175 mutations destabilize the CTC (Figure 6B). In contrast to destabilizing the isolated CTC, these very same ALS-disposing mutations significantly enhance the propensity of the intact LC domain of FUS to form runaway polymers (Figure 7A). Following the mutual exclusion teaching of correlate 2, variants of the FUS LC domain lacking a functional CTC are unable to deploy CTC-specified cross-β interactions to interfere with NTC interactions. As such, runaway NTC polymerization takes place (Figure 7A). We make note of the fact that this behavior of the G156E mutation within the FUS LC domain has already been documented by the groups of Hyman/Alberti and St George-Hyslop (24, 25).

We offer the schematic diagram shown in Figure 8C as an illustration of the concept of mutual exclusion. Kinetic parameters may favor initial self-association of the FUS LC domain to form the CTC cross-β structure (Figure 5B). The resulting proximity of two, unstructured NTC regions is predicted to facilitate a process of isomerization that begets formation of the slightly more stable NTC cross-β structure. Important to the thesis articulated herein, the concept of mutual exclusion demands that the initial CTC cross-β structure be disassembled in order for the NTC structure to form.

Once equilibrium has been reached, we offer two reasons to account for impediments to further growth of NTC polymers. First, if a third FUS LC domain were to use its CTC to invade the free CTC of the dimer held together by NTC cross-β interactions, mutual exclusivity would demand that NTC interactions dissolve (correlate 2). The same idea would guard against the ability of a third FUS LC domain to invade the free and unstructured NTC of a dimer held together by CTC cross-β interactions. Should run-away polymerization by either cross-β core take place, the correlate of mutual exclusion may no longer apply. If a series of NTC’s are co-assembled into a long polymer, we propose that an incoming monomer is unable to use its CTC to productively coalesce with any of the unstructured CTC domains extending laterally from the polymer. We offer that this option is voided because each NTC internal to the polymer is bordered on both sides by another structured NTC. Whereas the forces of mutual exclusion are proposed to be sufficient to force disassembly of a pair of molecules organized in the cross-β conformation (Figure 8C), they may be inadequate to force dissolution of a cross-β assembly bearing the forces of structural order from both sides.

Our second reason for hypothesizing orderly limitation of FUS polymer growth derives from correlate 1. Upon encountering either of the dimers shown in Figure 8C, an unstructured FUS LC domain would prefer interaction with the region of FUS not already existing in the cross-β structural state. If the dimer were held together by NTC interactions, the incoming protein is predicted to use its CTC in an attempt to form cross-β interactions with the unstructured CTC of the dimer. Reciprocally, if the dimer were held together by CTC interactions, the incoming protein would be expected to use its NTC to attempt self-association with the unstructured NTC of the dimer. These preferences, as specified in correlate 1, are attributed to conformal restriction. If the bias of conformational restriction is strong, and if mutual exclusivity is likewise strong, the combination of these limitations should prevent the FUS LC domain from polymerizing any further than the dimeric state.

We close with emphasis on the multitude of unknowns that cloud our simplistic ideas. The FUS LC domain is subject to many forms of post-translational modification (PTM) that may influence behavior of the NTC and CTC domains. Differential regulatory effects of PTMs might allow the FUS LC domain to expand either NTC- or CTC-mediated polymerization on demand. The FUS protein likewise shuttles from nucleus to cytoplasm and back, and uses its LC domain for any of a number of heterotypic interactions with other proteins. These variables can be understood to open the opportunity for the FUS LC domain to deviate from the dimeric ground state imposed by correlates 1 and 2. Despite the complexities of this science, we have faith in the value of the reductionist approach exemplified by the experiments described herein.

## Materials and Methods

### Cloning

Expression plasmids for mCherry:FUS LC, GFP:FUS LC and His-FUS LC were constructed as described previously (7). PCR fragments for preparing systematically truncated mutants of FUS LC domain, including ΔN8 and ΔC6, were amplified from full-length FUS LC domain as a template and sub-cloned into the multiple cloning site of the pHis-parallel1-GFP vector (27, 28). PCR fragments of ΔN8, ΔC6, FUS111-160 and FUS141-214 were also sub-cloned into the multiple cloning site of the pHis-parallel1-mCherry and pHis-parallel1-GFP vectors (27, 28). Caspase-3 cleavage site (GDEVD/A) was then inserted between GFP and the FUS fragments, resulting pHis-parallel1-GFP-GDEVDA-FUS fragments for production of tag-free proteins. PCR fragments ΔN8, ΔC6, FUS111-160 and FUS141-214 were also sub-cloned in pHis-parallel1 vector. PCR fragments corresponding to systematically deleted mutants of FUS141-214 were sub-cloned into the multiple cloning site of the pHis-parallel1-GFP-vector. ALS mutations in GFP-FUS LC, His-FUS LC, GFP-GDEVDA-ΔN8 and His-ΔC6 were introduced by QuikChange site-directed mutagenesis. All expression plasmids were confirmed by DNA sequencing.

### Protein expression and purification

All recombinant proteins were over-expressed in *E. coli* BL21(DE3) cells. Expression and purification of all GFP- or mCherry-fusion proteins were carried out as described previously (7, 28). Briefly, fusion proteins were overexpressed by induction with 0.5 mM IPTG at 20°C overnight. Cells were harvested and resuspended in lysis buffer containing 50 mM Tris-HCl pH 7.5, 500 mM NaCl, 10 mM β-mercaptoethanol (BME), 2 M urea and one tablet of protease inhibitor cocktail (Sigma). Cell suspensions were lysed by sonication for 2 minutes (10 seconds on/30 seconds off) on ice. Cell lysates were centrifuged at 142,000 × g for 1 hour at 4°C. Supernatants were mixed with Ni-NTA resin (Gold Bio), shaken gently for 15 min in a cold room, then poured into a glass column. The resin was washed with buffer containing 20 mM Tris-HCl pH7.5, 500 mM NaCl, 10 mM BME, 0.1 mM phenylmethylsulfonyl fluoride (PMSF), 2 M urea and 20 mM imidazole. Bound protein was eluted from the resin with an elution buffer containing 20 mM Tris-HCl pH7.5, 500 mM NaCl, 10 mM BME, 2 M urea and 250 mM imidazole. Purified proteins were concentrated using Amicon Ultra filters (Millipore). GFP fusion proteins were stored at −20°C as 50% glycerol stock and mCherry fusion proteins were stored at −80°C. Protein purity was confirmed by SDS-PAGE and concentrations were determined by absorbance at UV280.

His-FUS LC, His-ΔC6 and His-ΔN8 proteins were expressed and purified as described previously (8). Briefly, proteins were overexpressed by induction with 0.5 mM IPTG at 23°C overnight. Cells were harvested and resuspended in lysis buffer containing 50 mM Tris-HCl pH 7.5, 500 mM NaCl, 10 mM BME, 2 M guanidine hydrochloride (GdnHCl) and one tablet of protease inhibitor cocktail (Sigma). Cell suspension was lysed by sonication for 3 minutes (10 seconds on/30 seconds off) on ice. Cell lysates were centrifuged at 142,000 × g for 1 hour at 4°C. Supernatants were mixed with Ni-NTA resin (Gold Bio), shaken gently for 15 min in a cold room, then poured into a glass column. The resin was washed with buffer containing 20 mM Tris-HCl pH7.5, 500 mM NaCl, 10 mM BME, 2 M GdnHCl and 20 mM imidazole. Bound protein was eluted from the resin with an elution buffer containing 20 mM Tris-HCl pH7.5, 500 mM NaCl, 10 mM BME, 2 M GdnHCl and 250 mM imidazole. Purified proteins were concentrated using Amicon Ultra filters (Millipore). GdnHCl powder was added to the protein solution to make the final GdnHCl concentration 6 M. Protein solutions were stored at −80°C. Protein purity was confirmed by SDS-PAGE and the concentrations were determined by absorbance at UV_280_.

Tag-free FUS LC, dN8,ΔC6, FUS111-160, FUS141-214 and their point or deletion mutants were produced by cleavage of GFP tag with caspase-3 protease from His-GFP fusion proteins. Briefly, the concentrated protein was diluted to 2-3 mg/ml in a gelation buffer containing 20 mM Tris-HCl pH 7.5, 200 mM NaCl, 0.5 mM EDTA, 0.1 mM PMSF and 20 mM BME. Caspase-3 protease was then added at a 1:2000 ratio of the target protein. The reaction mixture was gently rotated at room temperature (RT) overnight. Cleaved proteins were mostly precipitated during the cleavage reaction and recovered by centrifugation at 4000 rpm for 30 min at 4°C. The precipitated proteins were dissolved in 2 ml of a denaturing buffer containing 20 mM Tris-HCl pH7.5, 200 mM NaCl, 10 mM β-ME, 0.5 mM EDTA, 0.1 mM PMSF and 6 M GdnHCl, and loaded onto a Superdex S-200 gel filtration column equilibrated with 20 mM Tris-HCl pH7.5, 200 mM NaCl, 10 mM β-ME, 0.5 mM EDTA and 2 M GdnHCl. Fractions containing the cleaved proteins were combined and concentrated to ~60 mg/ml. Powder of GdnHCl was added to the protein solution to make the final GdnHCl concentration 6 M, and samples were stored at −80 °C.

### Hydrogel binding assays

Hydrogel droplets of mCherry:FUS LC domain, mCherry:ΔN8 and mCherry:ΔC6 were prepared as described previously (28). GFP-test proteins were diluted in 2 ml of a gelation buffer containing 20 mM Tris-HCl pH7.5, 200 mM NaCl, 20 mM β-ME, 0.5 mM EDTA and 0.1 mM phenylmethylsulfonyl fluoride (PMSF) at the final concentration of 0.5 μM. Diluted protein solutions were applied onto hydrogel droplets in a 3.5-mm glass-bottomed cell culture dish (Mattek). After incubation at 4°C for overnight, the hydrogels were visually analyzed by fluorescence microscopy (Zeiss LSM880 Airyscan) as described previously (28).

### Polymer extension assays

Cross-β polymers of mCherry:FUS LC were prepared as described before (1). The polymer solution (~2 μM monomer concentration) was sonicated briefly to make polymer fragments. To this solution, GFP-FUS LC monomer solution was added at a 100 μM final concentration. After 10 min incubation at RT, the mixture was diluted 100 fold and the diluted polymer solution (10 μl) was placed on a coverslip (22 mm × 55 mm, 1.5 mm thickness), then a second coverslip (22 mm × 22 mm) was placed on the droplet. The polymers were visualized by Olympus total internal reflection fluorescent (TIRF) microscope with a 100x objective lens. Fluorescent images for the mCherry and GFP signals were recorded with a Hamamatsu CCD camera. Images were analyzed by ImageJ.

### Polymer formation

Tag-free FUS LC, ΔN8, ΔC6, FUS111-160, FUS141-214 and their point or deletion mutants in 6 M GdnHCl stock solution were diluted in 300 μl of the denaturing buffer at a final protein concentration of 2 mg/ml (FUS LC domain), 3 mg/ml (ΔN8 and ΔC6), or 5 mg/ml (FUS111-160 and FUS141-214) and then dialyzed against the gelation buffer for overnight. Protein solutions were recovered in microcentrifuge tubes. 10% sodium azide solution was added to a final concentration of 0.02%. Dialyzed solutions were incubated at room temperature for 1-2 weeks. On every other day of the first week, brief sonication was applied to facilitate polymerization.

### X-Ray diffraction

For X-ray diffraction, polymers of tag-free ΔN8, ΔC6, FUS111-160 and FUS141-214 were collected by centrifugation at 20,000 x g for 30 min. Polymer pellets were resuspended in 200 μl milli-Q water and dialyzed against 1 L milli-Q water for 12 hours twice. Dialyzed samples were lyophilized overnight and exposed to an X-ray beam to obtain diffraction patterns as described previously (7).

### Semi-denaturing detergent agarose gel electrophoresis (SDD-AGE)

Polymers of His-tag ΔN8, ΔC6, FUS111-160 and FUS141-214 or the yeast Sup35NM protein were diluted in the gelation buffer at 0.2 mg/mL (ΔN8, ΔC6, FUS111-160 and FUS141-214) or 0.1 mg/mL (Sup35NM), respectively, and sonicated briefly. Polymer solutions were incubated for 15-30 minutes in gelation buffer with indicated concentrations of SDS (0 - 2%) at 37 °C. Reaction mixtures were then loaded onto a 1.5% agarose gel to separate polymers and monomers. Proteins were transferred onto a nitrocellulose membrane and analyzed by western blotting using an anti-His tag antibody (7) or stained with Krypton fluorescent dye (ThermoFisher) according to a company’s instruction, and visualized by Typhoon FLA 9000 gel scanner (GE).

### Thioflavin-T assays

Tag-free FUS LC domain, ΔN8 or ΔC6 proteins, Fus111-160, FUS141-214 and their ALS variants were diluted to 25 μM (ΔN8 and ΔC6) or 50 μM (FUS LC domain) with 60 μL of a ThT buffer containing 20 mM Na-phosphate, pH 7.4, 100 mM NaCl, 0.5 mM EDTA, 0.1 mM PMSF, 1 mM TCEP and 40 μM Thioflavin-T. Diluted samples were transferred to a 96 well fluorescence assay plate (Costar assay plate 96, CORNING). The plate was sealed with a foil film to prevent evaporation. Thioflavin-T signal was monitored with a CYTATION5 plate reader using an excitation wavelength at 450 nm and an emission wavelength at 490 nm. Reading was carried out every 15 minutes with 15 seconds shaking being applied before each reading. The ThT buffer alone was used as a reading blank. Experiments were performed in 3-6 multiplicate.

### Thermal stability assays

Cross-β polymers of ΔN8 or ΔC6 were prepared as described above. The polymer solutions were ultracentrifuged at 75,000 × g for 1 hour. Supernatants were removed, and 300 μl of fresh gelation buffer was added to the polymers. Brief sonication was applied to resuspend the polymer pellets, then the polymers were again ultracentrifuged at 75,000 × g for 1 hour. Supernatants were removed, and 300 μl of fresh gelation buffer was added. Brief sonication was again applied to resuspend the polymer pellets. The polymer solutions were incubated 30°C for 1 hour, then ultracentrifuged at 75,000 × g for 1 hour. Supernatants were removed, and 300 μl of fresh gelation buffer was added. Brief sonication was again applied to resuspend the polymer pellets. The polymer solutions were incubated 40°C for 1 hour, then ultracentrifuged at 75,000 x g for 1 hour. These steps were repeated with a 10°C increment of the incubation temperature until 70°C incubation was finished. The supernatants from each temperature step (300 μl each) were mixed with 700 μl of RPC buffer A containing 2% acetonitrile and 0.065% trifluoroacetic acid. The pellet of the 70°C incubation was resuspended in the mixture of 300 μL of gelation buffer and 700 μl of RPC buffer A. Release of monomers of ΔN8 or ΔC6 in these samples was analyzed by reverse phase column chromatography (Resource RPC column: Cytiva) with 0 to 60% acetonitrile gradient in 0.05% trifluoroacetic acid. Peak areas for ΔN8 or ΔC6 were recorded, converted to protein concentrations, then plotted against temperature. Experiments were performed in triplicate.

### Sample preparation for ssNMR

To prepare ^13^C, ^15^N labeled tag-free ΔN8, ΔC6, FUS111-160 and FUS141-214, proteins were expressed in BL21(DE3) *E. coli* cells cultured in M9 media with ^13^C-glucose as a carbon source and ^15^N-ammonium chloride as a nitrogen source at 20°C for overnight. Proteins were purified as described above. The protein fragments were cleaved from the His-GFP tag by caspase-3 and then further purified with an S-200 Superdex gel filtration column S-200. Fractions containing object proteins were combined and concentrated. To make fibril samples, proteins were diluted in 6 M guanidine hydrochloride at 3 mg/ml (ΔN8 and ΔC6) or 5 mg/ml (FUS111-160 and FUS141-214), and then dialyzed in a buffer containing 20 mM Na-phoshpate buffer pH 7.4, 20 mM NaCl, 20 mM β-ME, 0.5 mM EDTA, 0.1 mM PMSF and 0.02% Na azide at 25°C overnight. The dialyzed proteins were recovered and incubated at 4 C° for two weeks. To facilitate polymer formation, the protein solution was sonicated several times during the incubation.

For segmental labeling of His-GFP-FUS LC, an artificially designed Cfa intein was employed as described previously (8, 13). Briefly, a N-terminal construct His-GFP-GG-Cfa was expressed in LB medium at 20°C overnight. A C-terminal construct His-GFP-GDEVDC-FUS LC was expressed in M9 minimal media with ^13^C-glucose and ^15^N-ammonium chloride at 20°C for overnight. Purification of the fusion constructs was carried out as described before (8, 13). Isotopically-labeled His-GFP-GDEVDC-FUS LC and unlabeled His-GFP-GG-Cfa were mixed at a 1:1 molar ratio (100 μM: 100 μM) in a ligation buffer containing 20 mM Na-phosphate buffer (pH 7.4), 150 mM NaCl, 1 mM TCEP, 400 mM MESNa and 30 μg/mL caspase-3 at 25°C for 48 hours. The ligation products were recovered by centrifugation and resolved in a buffer containing 20 mM Tris-HCl (pH 7.5), 200 mM NaCl, 20 mM β-ME and 4 M urea. The solubilized products were concentrated and loaded on an S-200 gel filtration column (Cytiva, USA) equilibrated with a buffer containing 20 mM Tris-HCl (pH 7.5), 200 mM NaCl, 20 mM β-ME and 2 M urea. Fractions containing the segmentally-labeled His-GFP-GGC-FUS LC were combined, concentrated and dialyzed against a buffer containing 20 mM Na-phosphate buffer (pH 7.4), 20 mM NaCl, 20 mM β-ME, 0.5 mM EDTA, 0.1 mM PMSF and 0.02% Na azide at 25°C overnight. The dialyzed proteins were recovered and incubated at 4 C° for two weeks. To facilitate polymer formation, the protein solution was sonicated several times during the incubation.

Fibril samples were collected and packed in a thin-wall 3.2-mm nmr rotor by centrifugation with a homemade packing device. Data were acquired at 14.1 T with a Tecmag Redstone spectrometer console and a Varian 3.2 mm BioMAS probe. MAS frequencies were 12 kHz. Sample temperatures were 30°C, based on the ^1^H NMR frequency of water in the samples. 2D ^13^C-^13^C spectra used 70 kHz TPPM decoupling in t1 and t2 dimensions. 2D ^13^C-^13^C spectra used 25 ms of RAD/DARR mixing, 200 complex t1 points with a 20 microsecond t1 increment, and 80-256 scans per FID with a 1.5 second recycle delay. ^1^H and ^13^C RF fields were 55 kHz and 43 kHz during ^1^H-^13^C cross-polarization, with a 1.5 ms contact time. 2D data were processed in nmrPipe (29) with 0.67 ppm Gaussian line broadening in ^13^C dimension.

## Acknowledgements

We thank Robert Tycko and Myungwoon Lee for help in recording ssNMR spectra. We thank Deepak Nijhawan and Glen Liszczak for thoughtful discussions regarding the research described herein. We also thank Lillian Sutherland, Lily Sumrow and Leeju Wu for plasmid cloning. SLM was supported by NIGMS grant 5R35GM130358 as well as unrestricted funding from an anonymous donor. Experiments were planned by MK and SLM. Biochemical and molecular biological experiments were performed by MK. MK and SLM composed the manuscript. Neither author of this manuscript has any form of conflict or competing interest. All data in this manuscript and supplementary materials will be freely available upon publication.

## Supplemental Data Figure Legends

**Figure S1.**
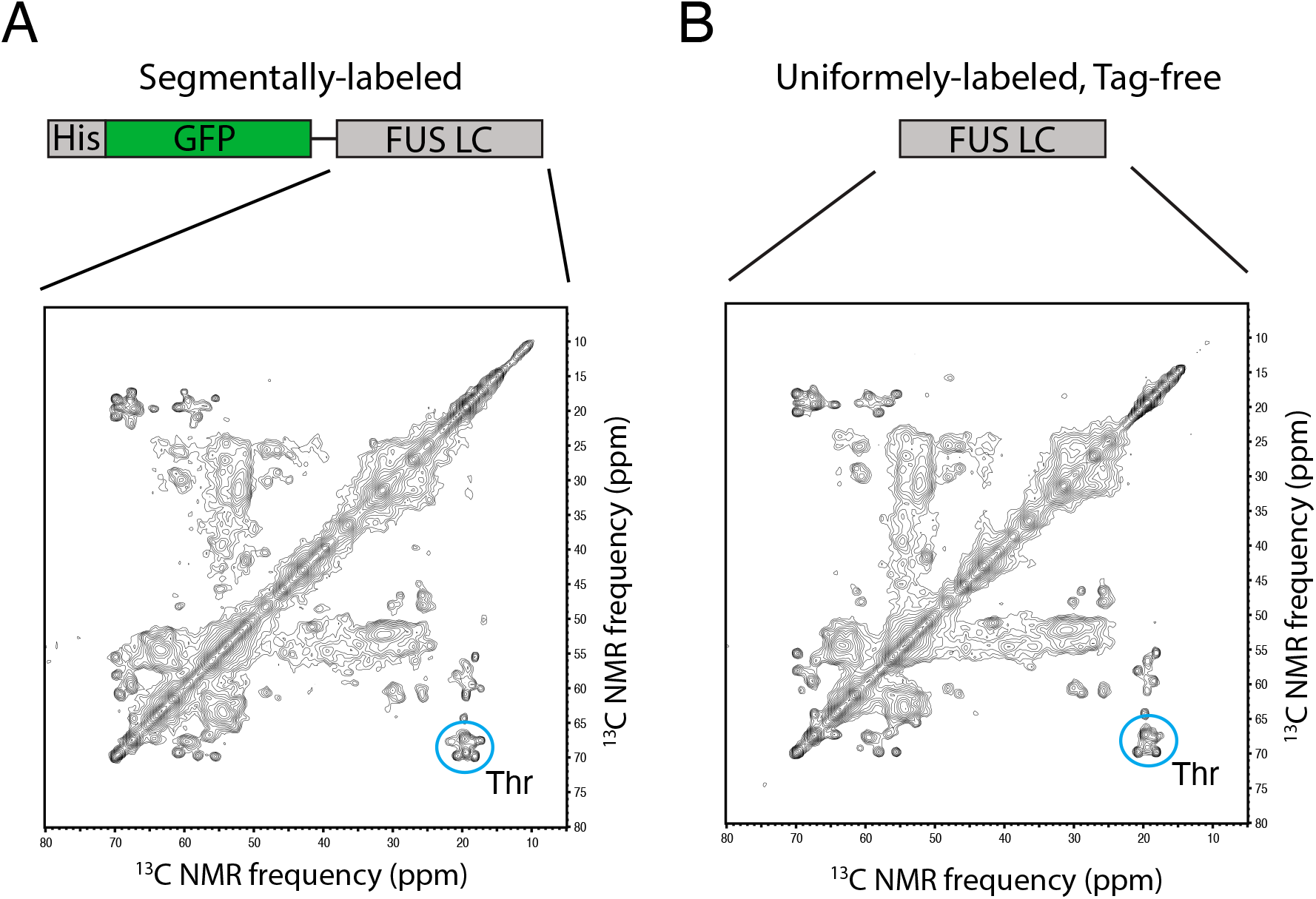
Solid state NMR spectra of segmentally labeled GFP-FUS polymers relative to polymers made from tag-free LC domain of FUS. (A) 2D ^13^C-^13^C spectra of segmentally labeled GFP-FUS LC polymers. The segmentally-labeled protein was prepared using intein chemistry, purified and incubated under conditions receptive to polymerization (Materials and Methods). (B) 2D ^13^C-^13^C spectra of uniformly labeled tag-free FUS LC polymers. The tag-free protein was prepared by cleaving His-tag from His-tag FUS LC with TEV, purified and incubated under conditions receptive to polymerization (Materials and Methods). The spectral peaks circled in blue represent threonine residues that are exclusively found in the N-terminal core of FUS LC (no threonine residue in the C-terminal half). Contour levels increase by successive factors of 1.30 in both the panels.

**Figure S2.**
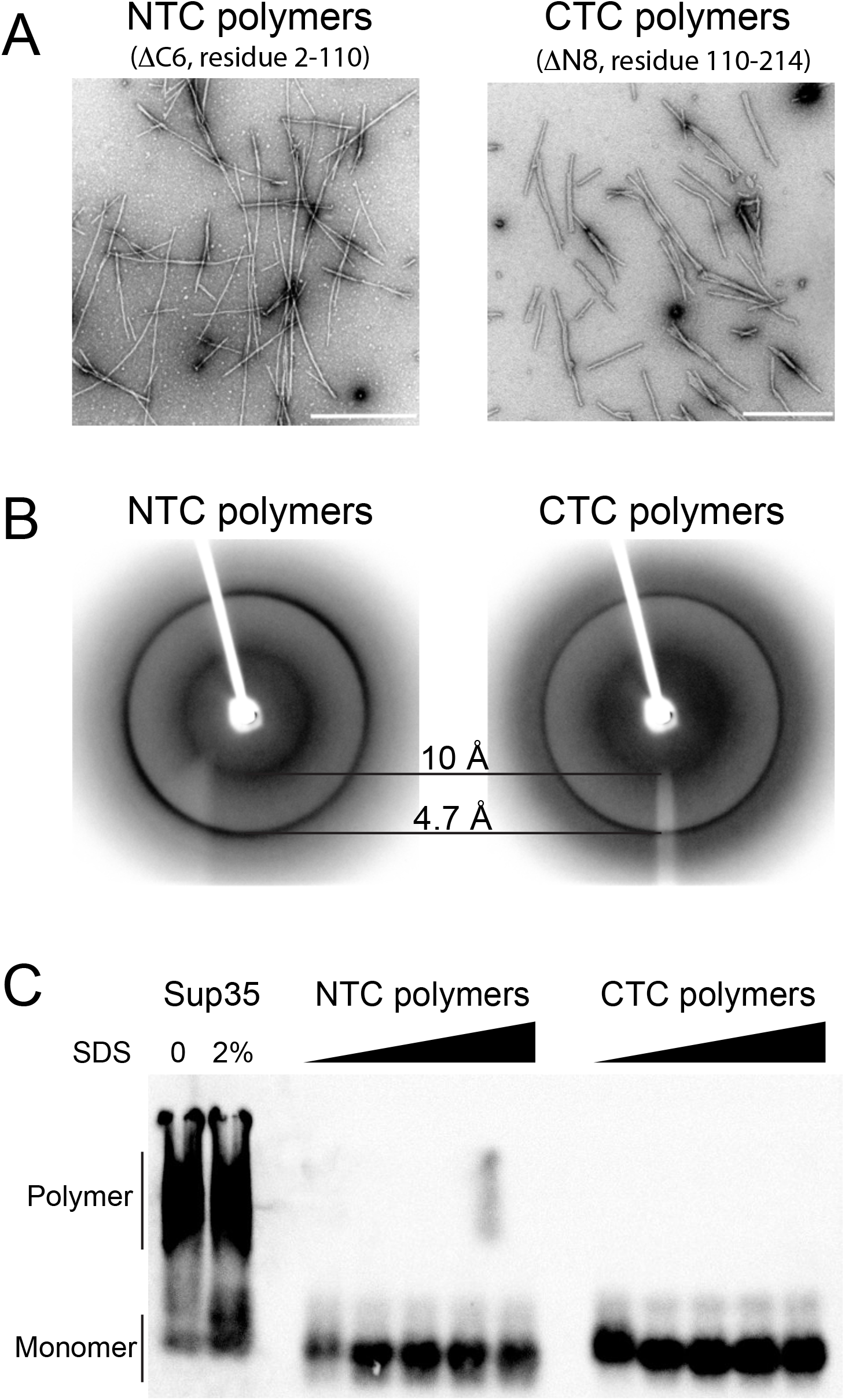
Properties of cross-β polymers formed from the C-terminal half of the FUS LC domain. (A) The ΔN8 deletion variant (Figure 2) spans residues 111-214 of the FUS LC domain. Upon incubation at neutral pH and physiological concentrations of monovalent salt, this protein fragment polymerized into homogenous, unbranched polymers as observed by transmission electron microscopy. (B) A lyophilized hydrogel droplet containing these polymers was irradiated with X-rays revealing prominent diffraction rings at 4.7 Å and 10 Å. (C) Unlike irreversible amyloid polymers formed from the yeast Sup35 protein (left side of panel), polymers formed from the ΔN8 variant dissolved to the monomeric state upon assay by semi-denaturing agarose gel electrophoresis (right side of panel).

**Figure S3.**
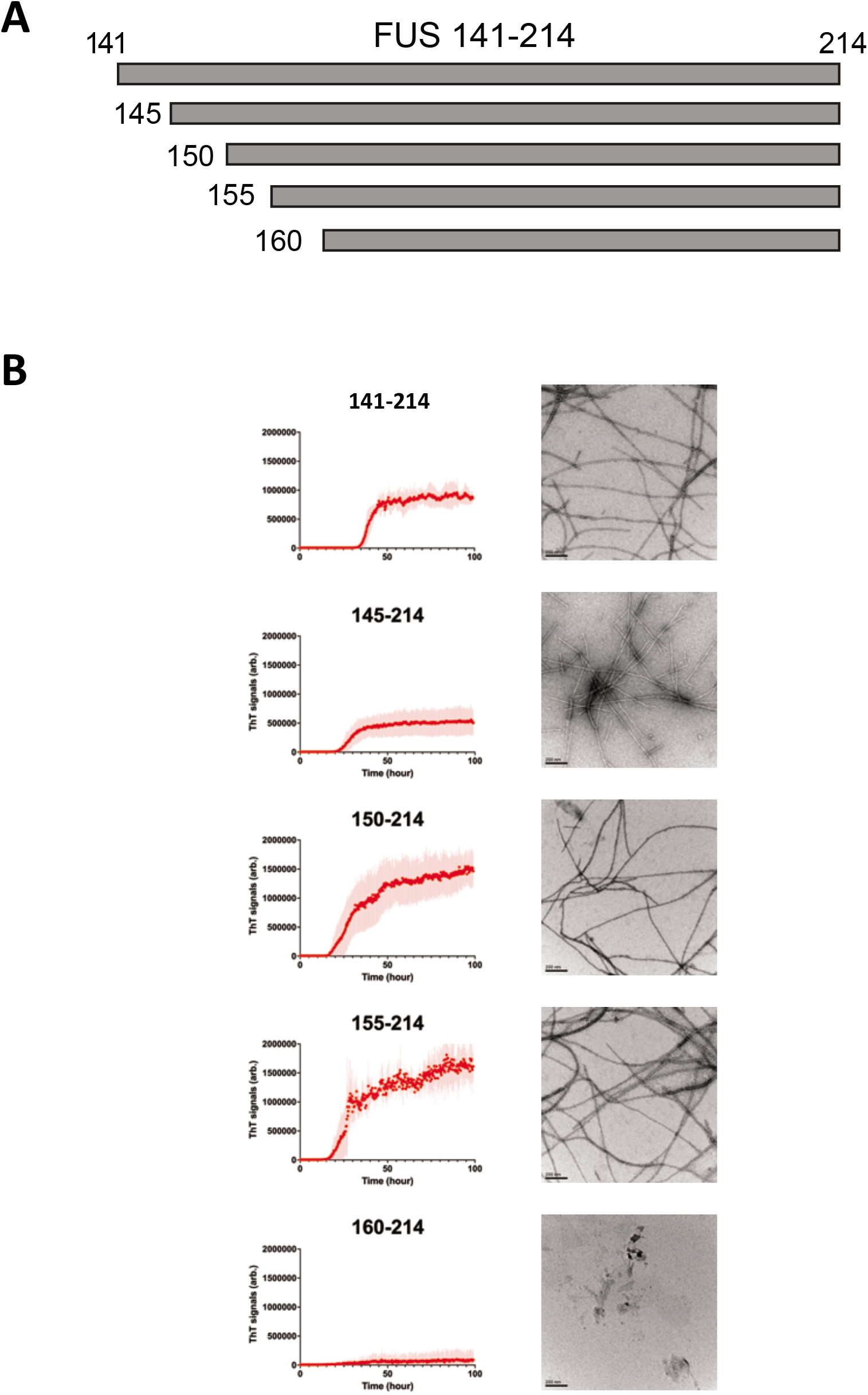
Polymerization assays for truncated variants of C-terminal half of the FUS LC domain. (A) Segments of the FUS LC domain containing N-termini at residues 141, 145, 150, 155 and 160, and commonly terminating at residue 214 were purified and incubated under conditions receptive to polymerization (Materials and Methods). (B) Assays for time-dependent acquisition of thioflavin-T fluorescence (left) and electron microscopy (right) were used to monitor the formation of amyloid-like polymers. Scale bar = 200 μM.

**Figure S4.**
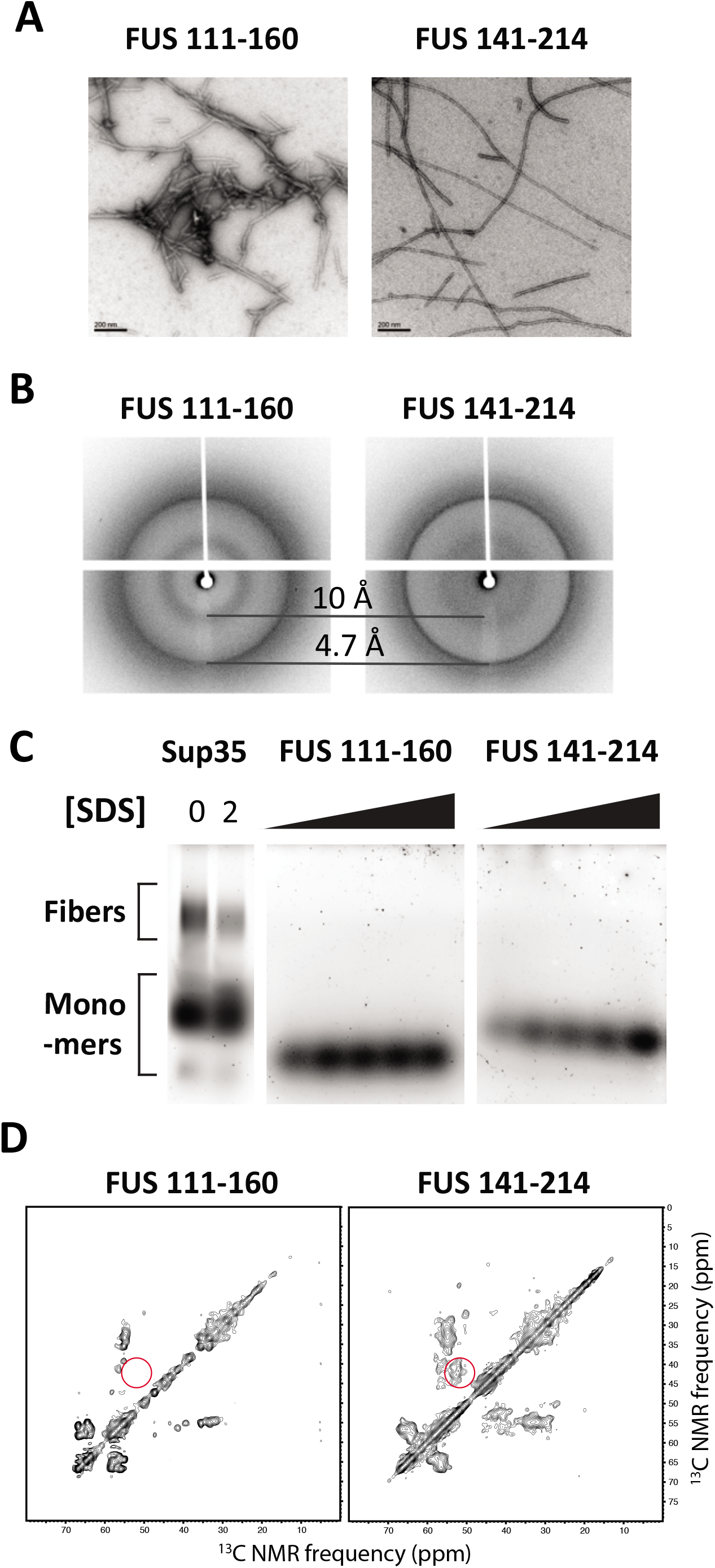
Electron microscopy, X-ray diffraction, semi-denaturing agarose gel electrophoresis, and solid state NMR analysis of cross-β cores made from two distinct regions of the C-terminal half of the FUS LC domain. Purified fragments spanning either residues 111-160 or 141-214 of the FUS LC domain were incubated under conditions receptive to formation of cross-β polymers. (A) Electron microscopic images of polymers made from 111-160 fragment (left) or 141-214 fragment (right) of FUS LC domain. (B) X-ray diffraction images of lyophilized polymer samples of 111-160 fragment (left) or 141-214 fragment (right). (C) Semidenaturing agarose gel electrophoresis analysis of polymers made from yeast Sup35 protein (left), 111-160 fragment (middle), or 141-214 fragment (right) of FUS LC domain. (D) 2D ^13^C-^13^C spectra of uniformly labeled polymers made from 111-160 fragment (left) or 141-214 fragment (right). Red cercles indicate a presence of rigid Asn/Asp residues in the core structure of 141-214 fragment, but not in 111-160 fragment. Contour levels increase by successive factors of 1.3 in both the panels.

## References

1. N. Rado-Trilla, M. Alba, Dissecting the role of low-complexity regions in the evolution of vertebrate proteins. BMC Evol Biol 12, 155 (2012).

2. M. Toll-Riera, N. Rado-Trilla, F. Martys, M. M. Alba, Role of low-complexity sequences in the formation of novel protein coding sequences. Mol Biol Evol 29, 883–886 (2012).

3. P. Leuenberger et al., Cell-wide analysis of protein thermal unfolding reveals determinants of thermostability. Science 355(2017).

4. S. F. Banani, H. O. Lee, A. A. Hyman, M. K. Rosen, Biomolecular condensates: organizers of cellular biochemistry. Nat Rev Mol Cell Biol 18, 285–298 (2017).

5. Y. Shin, C. P. Brangwynne, Liquid phase condensation in cell physiology and disease. Science 357(2017).

6. T. W. Han et al., Cell-free Formation of RNA Granules: Bound RNAs Identify Features and Components of Cellular Assemblies. Cell 149, 768–779 (2012).

7. M. Kato et al., Cell-free Formation of RNA Granules: Low Complexity Sequence Domains Form Dynamic Fibers within Hydrogels. Cell 149, 753–767 (2012).

8. D. T. Murray et al., Structure of FUS Protein Fibrils and Its Relevance to Self-Assembly and Phase Separation of Low-Complexity Domains. Cell 171, 615–627 e616 (2017).

9. R. Tycko, Solid-state NMR studies of amyloid fibril structure. Annu Rev Phys Chem 62, 279–299 (2011).

10. M. Fändrich, J. Meinhardt, N. Grigorieff, Structural polymorphism of Alzheimer Aβ and other amyloid fibrils. Prion 3, 89–93 (2009).

11. S. Xiang et al., The LC Domain of hnRNPA2 Adopts Similar Conformations in Hydrogel Polymers, Liquid-like Droplets, and Nuclei. Cell 163, 829–839 (2015).

12. V. H. Ryan et al., Mechanistic View of hnRNPA2 Low-Complexity Domain Structure, Interactions, and Phase Separation Altered by Mutation and Arginine Methylation. Mol Cell 69, 465–479 e467 (2018).

13. D. T. Murray et al., Structural characterization of the D290V mutation site in hnRNPA2 low-complexity-domain polymers. Proc Natl Acad Sci U S A 115, E9782–E9791 (2018).

14. J. Lu et al., CryoEM structure of the low-complexity domain of hnRNPA2 and its conversion to pathogenic amyloid. Nature Communications 11, 4090 (2020).

15. H. J. Kim et al., Mutations in prion-like domains in hnRNPA2B1 and hnRNPA1 cause multisystem proteinopathy and ALS. Nature 495, 467–473 (2013).

16. N. M. Vieira et al., A defect in the RNA-processing protein HNRPDL causes limb-girdle muscular dystrophy 1G (LGMD1G). Hum Mol Genet 23, 4103–4110 (2014).

17. Y. Lin et al., Toxic PR Poly-Dipeptides Encoded by the C9orf72 Repeat Expansion Target LC Domain Polymers. Cell 167, 789–802 e712 (2016).

18. N. Ticozzi et al., Analysis of FUS gene mutation in familial amyotrophic lateral sclerosis within an Italian cohort. Neurology 73, 1180–1185 (2009).

19. T. J. Kwiatkowski, Jr. et al., Mutations in the FUS/TLS gene on chromosome 16 cause familial amyotrophic lateral sclerosis. Science 323, 1205–1208 (2009).

20. R. Rademakers et al., Fus gene mutations in familial and sporadic amyotrophic lateral sclerosis. Muscle & nerve 42, 170–176 (2010).

21. M. Lee, U. Ghosh, K. R. Thurber, M. Kato, R. Tycko, Molecular structure and interactions within amyloid-like fibrils formed by a low-complexity protein sequence from FUS. Nat Commun 11, 5735 (2020).

22. T. J. Kwiatkowski et al., Mutations in the FUS/TLS Gene on Chromosome 16 Cause Familial Amyotrophic Lateral Sclerosis. Science 323, 1205 (2009).

23. J. Yan et al., Frameshift and novel mutations in FUS in familial amyotrophic lateral sclerosis and ALS/dementia. Neurology 75, 807–814 (2010).

24. A. Patel et al., A Liquid-to-Solid Phase Transition of the ALS Protein FUS Accelerated by Disease Mutation. Cell 162, 1066–1077 (2015).

25. T. Murakami et al., ALS/FTD Mutation-Induced Phase Transition of FUS Liquid Droplets and Reversible Hydrogels into Irreversible Hydrogels Impairs RNP Granule Function. Neuron 88, 678–690 (2015).

26. R. M. Vernon et al., Pi-Pi contacts are an overlooked protein feature relevant to phase separation. Elife 7(2018).

27. P. Sheffield, S. Garrard, Z. Derewenda, Overcoming expression and purification problems of RhoGDI using a family of “parallel“ expression vectors. Protein expression and purification 15, 34–39 (1999).

28. M. Kato, Y. Lin, S. L. McKnight, Cross-beta polymerization and hydrogel formation by low-complexity sequence proteins. Methods 126, 3–11 (2017).

29. F. Delaglio et al., NMRPipe: A multidimensional spectral processing system based on UNIX pipes. Journal of Biomolecular NMR 6, 277–293 (1995).

